# The HIV-1 capsid core is an opportunistic nuclear import receptor

**DOI:** 10.1101/2021.12.02.470925

**Authors:** Guangai Xue, Hyun Jae Yu, Shih Lin Goh, Anna T. Gres, Mehmet Hakan Guney, Stefan G. Sarafianos, Jeremy Luban, Vineet N. KewalRamani

## Abstract

The movement of viruses and other large macromolecular cargo through nuclear pore complexes (NPCs) is poorly understood. The human immunodeficiency virus type 1 (HIV-1) provides an attractive model to interrogate this process due to the genetic and cell biological assays to score virus nuclear entry in living cells. Although initial studies of HIV-1 infection of nondividing cells focused on karyophilic virion proteins, subsequent work revealed the viral capsid (CA), the chief structural component of the pre-integration complex (PIC), to be a critical determinant in nuclear transport^1^. In support of this model, HIV-1 interactions with NPCs can be altered through CA mutation^2^, which makes direct contact with nucleoporins (Nups)^3–5^. Here we identify Nup35, Nup153, and POM121 to coordinately support HIV-1 nuclear entry. For Nup35 and POM121, this dependence was strongly dependent cyclophilin A (CypA) interaction with CA. Mutation of CA or removal of soluble host factors changed the interaction with the NPC. Collectively, these findings implicate the HIV-1 CA hexameric lattice that encapsulates the viral genome as a macromolecular nuclear transport receptor (NTR) that exploits soluble host factors to modulate NPC requirements during nuclear invasion.

---

HIV-1 exhibits significant flexibility in its utilization of the NPC to access the nucleus during early replication steps. Comprised of over 30 distinct proteins, the NPC incorporates six subcomplexes based on stoichiometry and distribution within the NPC (Extended Data Fig. 1a) ^6^. Approximately one-third of Nups contain FG-dipeptides, motifs which are targets for NTR interaction for directional movement of cargo from the cytoplasm to the nucleus ^7^. Genetic and biochemical studies have implicated the FG-Nups Nup153 and Nup358 as contributing to HIV-1 nuclear entry ^5,8–11^. These Nups decorate opposite poles of the NPC with tentacles of Nup358 extending into the cytoplasm from the outer nuclear membrane and Nup153 projecting a basket-like structure from the inner nuclear membrane into the nucleoplasm. HIV-1 with mutations in CA that prevent interaction with soluble factors such as CPSF6 and CypA, exhibit a differential dependence on Nups during infection, including loss of reliance on both Nup153 and Nup358. CypA interacts with CA via the P90 residue exposed on loop in the N-terminal domain (NTD), and CPSF6 interacts via the N74 pocket within an NTD and C-terminal domain (CTD) interface present in hexameric CA in the virion core (Extended Data Fig. 1b and 1c). P90A and N74D mutations, respectively, impair CypA and CPSF6 interactions with CA. We considered the possibility that CPSF6 and CypA binding to HIV-1 in the cell cytoplasm affected subsequent interactions at the NPC and thus routes of nuclear entry.

To better understand the applicability of this interface exposure model, we performed a genetic screen with small-interfering RNAs (siRNAs) targeting 32 human Nups in HeLa cells to identify those that exhibited CA-dependence during infection with VSV-G pseudotyped HIV-1 vectors encoding red fluorescent protein (RFP). As a positive control to inhibit infection of HIV-1 with wild-type (WT) CA, TNPO3 was also depleted using siRNA ^8^. Parallel infections were performed with HIV-1 CA mutant (N74D and P90A) virus vectors and Moloney murine leukemia virus (MLV) vectors. Nup35, Nup153, Nup358, and POM121 depletion were observed to inhibit WT HIV-1 infection but affected N74D and P90A HIV-1 infection to a lesser degree (Fig. 1a). Nup35 and POM121 have not previously been studied as potential HIV-1 cofactors, and notably, their depletion had little effect on MLV infection (Extended Data Fig. 2). As has been previously reported ^2,11^, Nup160 depletion affected WT and CA mutant HIV-1 infection, which is likely due to the critical role it plays in NPC formation.

**Figure 1.**
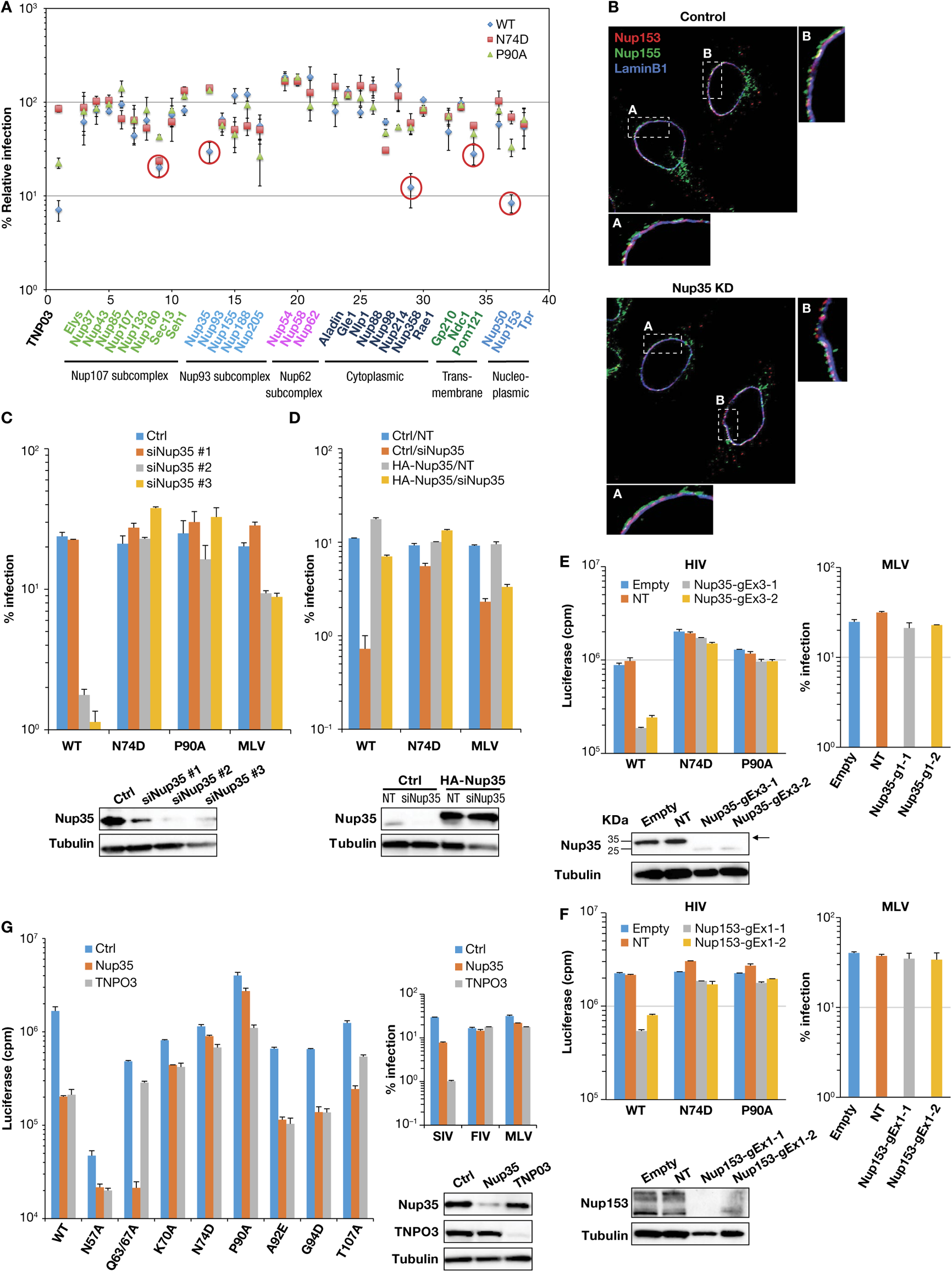
HIV-1 dependence on Nup35. **a,** CA-dependent use of nucleoporins by HIV-1. HeLa cells were transfected with a control siRNA or target siRNA (smart pool) for 48 h, then infected with m.o.i of 0.5 of VSV-G-pseudotyped wild-type (WT) virus or virus harboring various CA mutants. After 48 h, infected RFP-positive cells were counted by FACS. Nups targeted by siRNAs are grouped by NPC subcomplexes. TNPO3 siRNA-treated cells are included in all the screens as a positive control. WT HIV-1 infection decreases of greater than 3-fold after knockdown of nucleoporins are encircled. Percent relative infection is mean ± s.d., from three independent experiments. MLV infection was also measured at 48 h in parallel samples to monitor cell toxicity and specificity of HIV infection (Extended Data Fig. 2). **b**, Distribution of Nup153 and Nup155 after Nup35 knockdown. Localization of Nup153 (red) and Nup155 (green) in HeLa cells by deconvolution microscopy after immunostaining. Lamin B1 is blue. A Z-section image is presented. **c,** different siRNAs targeting Nup35 impair HIV-1 infection. HeLa cells were transfected with a control siRNA or three different siRNAs against Nup35 for 48 h, then infected with VSV-G-pseudotyped WT HIV-1 vectors or virus harboring various CA mutants or MLV. After 48 h, infected cells expressing fluorescent markers from viral vectors were enumerated by FACS. Error bars show standard deviations of duplicates, representative of three independent experiments. Western blot analysis confirmed Nup35 depletion by three different siRNAs. **d,** HeLa cells stably expressing human HA-Nup35 lacking 3’-UTR or transduced with control vector (LPCX-HA), were transfected with non-target (NT) siRNA or Nup35 siRNA targeting 3’-UTR region for 48 h, then infected with VSV-G-pseudotyped virus. Infected RFP-positive cells were counted by FACS. Error bars show standard deviations of duplicates, representative of three independent experiments. Western blot confirmed Nup35 restoration. **e**, **f**, HIV-1 CA determines Nup35 (**e**) or Nup153 (**f**) dependence in HeLa knockout cell clones. Nup35 or Nup153 knockout HeLa cell clones were infected by HIV-1 or MLV. Infected RFP-positive cells were counted by FACS 48 h after infection. Error bars show standard deviations of duplicates, representative of three independent experiments. Western blots show the knockout efficiency of Nup35 (**e**) or Nup153 (**f**). Empty, cell lines that only transiently co-transfected pX330 with puromycin-expression vector, plasmid; NT, non-targeting control lines lacking a gene-specific gRNA. **g**, HeLa cells were transfected with control siRNA or siRNAs against Nup35 or TNPO3 for 48 h, then infected with VSV-G-pseudotyped pNL4-3-Luc-E-R+ or virus harboring various CA mutants, or various retroviruses as labeled. Firefly luciferase activity and GFP/RFP-positive cells were monitored at 48h. Error bars show standard deviations of duplicates, representative of two independent experiments. Western blotting confirmed knockdown efficiency.

By contrast, depletion of Nup62 or Nup214 elevated WT HIV-1 infection, and Nup54 and Nup58 knockdown enhanced infection of both WT and CA mutant HIV-1. MLV infection was not increased in Nup54, Nup58, Nup62, or Nup214 knockdown cells (Extended Data Fig. 2). Nup54, Nup58, Nup62, and Nup214 are notable due to their formation of an “FG-hydrogel” that provides a selective barrier to transport between cytoplasm and nucleus.

Because Nup35 and POM121 depletion affected WT HIV-1 infection but not CA mutant HIV-1 in a pattern similar to Nup153 and Nup358, we focused efforts on these two Nups. Initial studies were performed with Nup35 as it was less essential to N74D and P90A HIV-1 relative to POM121. Nup153 was selected over Nup358 as a control in subsequent experiments given significant cellular toxicity observed after Nup358 knockdowns. Nup153 was also of interest because HIV-1 CA engages it via an FG-motif.

To determine whether the effect of Nup35 on WT HIV-1 infection was direct, we sought to examine the stability of other Nups after Nup35 depletion with siRNA. Knockdown of Nup35 did not affect levels of other Nups present in the Nup93 subcomplex of which it is a member nor did it affect the levels of TNPO3, Nup153, or Nup358 (Extended Data Fig. 3). These data suggested that the indirect degradation of other Nups, including those known to interact with HIV-1, did not account for the infection block. In addition, Nup153 continues to associate with the nuclear membrane in Nup35 knockdown cells as does the Nup93 subcomplex constituent Nup155, in the same relative orientation to one another (Fig. 1b). Because N74D and P90A HIV-1 infection are unimpeded in Nup35-depleted cells, coupled with the microscopy data, these results indicate transient Nup35 loss did not grossly affect nuclear transport or NPC integrity.

We next tested three unique Nup35 siRNAs from the mixture used in the screen separately. Consistent with the pooled siRNA results, two different siRNAs directed against Nup35, siRNA-2 and siRNA-3, reduced WT HIV-1 infection approximately 10-fold (Fig. 1c) correlating with Nup35 depletion, whereas N74D HIV-1 and P90A HIV-1 were resistant to Nup35 depletion.

To test whether the reductions in HIV-1 infection were due to off-target effects, we performed a functional rescue experiment (Fig. 1d). Nup35 siRNA-3 targets 3’-UTR sequence. Thus HeLa cells were transduced either with LPCX empty vector or with LPCX vector expressing HA-tagged Nup35 that lacks the 3’-UTR and is therefore resistant to silencing by the siRNA. Western blot analysis confirmed that siRNA knockdown reduced the endogenous level of Nup35 in both transduced cell populations; however, ectopically expressed HA-Nup35 was unaffected by Nup35 siRNA treatment. Consistent with prior results, cells depleted for endogenous Nup35 and transduced with the LPCX empty vector were more than 10-fold less susceptible to HIV-1 infection. In contrast, Nup35 siRNA-treated cells that expressed HA-Nup35 remained susceptible to WT HIV-1. N74D HIV-1 infection was relatively consistent under the different Nup35 expression conditions. Notably, the small infection decrease in MLV infection occurring after Nup35 knockdown was not offset in cells expressing exogenous HA-Nup35 suggesting the effect on MLV could be indirect.

CRISPR/Cas9 gene-editing of *nup35* in HeLa cells confirmed the siRNA findings. While 5 different guide RNAs (gRNAs) were tested, only one targeting the third exon of *nup35* yielded cell clones deficient of full-length Nup35. These clones uniformly expressed a truncated form of Nup35 that likely initiated from an in-frame methionine at coding residue position 62 of Nup35 within exon 3 based on the protein size and sequencing analysis (Fig. 1e and Extended Data Fig. 4a). The internally initiated form of Nup35 may have permitted cell survival. These cells expressing the suspected N-terminally truncated Nup35 had reduced susceptibility to HIV-1 infection but not N74D or P90A HIV-1 infection (Fig. 1e). Consistent with siRNA knockdown results, knockout of Nup35 did not affect levels of other Nups present in the Nup93 subcomplex nor did it affect the levels of TNPO3, Nup153, or Nup358 (Extended Data Fig. 4b). Knockout cell clones for Nup153 (Extended Data Fig. 4c) were also obtained and were similarly less permissive for HIV-1 infection relative to N74D or P90A HIV-1 infection (Fig. 1f). Although we were unable to detect a truncated form of Nup153 in the knockout cells by western blot analysis, it is possible that a shorter form permitting survival was not recognized by our antibodies.

Because N74D HIV-1 and P90A HIV-1 were insensitive to Nup35 depletion, we examined other viruses with CA mutations that influence interaction with CPSF6 or CypA for infection of Nup35 knockdown cells. HIV-1 with N57A, Q63A/Q67A, K70A, or T107A mutations in CA are reduced in sensitivity to CPSF6-mediated infection blocks and are similarly diminished in binding CPSF6 ^12^. Indeed, relative to WT HIV-1, these viruses were less sensitive to TNPO3 depletion which enables CPSF6 inhibition of infection (Fig. 1g). While N57A, K70A, and N74D HIV-1 were similarly less sensitive to Nup35 depletion, Q63A/Q67A and T107A HIV-1 resembled WT HIV-1 in infection indicating that CPSF6-interaction does not predict Nup35 dependence.

In contrast to WT HIV-1, A92E HIV-1 and G94D HIV-1 replicate more efficiently in the absence of CypA ^13,14^. They however retain CypA as well as CPSF6 interaction sites. Similar to WT HIV-1, these viruses were impaired for infection of Nup35 and TNPO3 knockdown cells (Fig. 1g). We similarly tested simian immunodeficiency virus from macaques (SIV_mac239_), which replicates independent of CypA interaction, and feline immunodeficiency virus (FIV) which does not interact with either CypA or CPSF6. SIV_mac239_, which retains the conserved region necessary for CPSF6 interaction, was potently impaired by TNPO3 knockdown but exhibited reduced dependence on Nup35 (Fig. 1g). FIV was unaffected by either Nup35 or TNPO3 knockdown. Taken together, these results demonstrate that HIV-1 CA determines sensitivity to Nup35 depletion, and the ability to interact with CypA correlated with dependence on Nup35.

## CypA impairs HIV-1 infection in Nup35 knockdown cells

To directly test whether CypA was required for the HIV-1 dependence on Nup35, we treated Nup35-knockdown cells with CsA to inhibit the CypA-CA interaction (Fig. 2a). WT HIV-1 infectivity was rescued to the level of control cells by CsA treatment in Nup35-knockdown cells. In contrast, CsA had minor effects on WT HIV-1 infection in either control cells or TNPO3-knockdown cells. N74D HIV-1 infection was previously demonstrated to be sensitive to CsA treatment ^15^, and it remained sensitive in Nup35- and TNPO3-knockdown cells. As expected, the infectivity of HIV-1 with the CypA binding mutation P90A was not affected by CsA treatment in any cell type. These data demonstrate that CsA restores HIV-1 replication following Nup35 knockdown. The effect was specific to HIV-1, as SIV_mac239_, FIV, and MLV were not affected by CsA in Nup35-depleted cells (Fig. 2a). Taken together, these data indicate that the block to HIV-1 infection in Nup35-depleted cells is dependent on the virus binding CypA.

**Figure 2.**
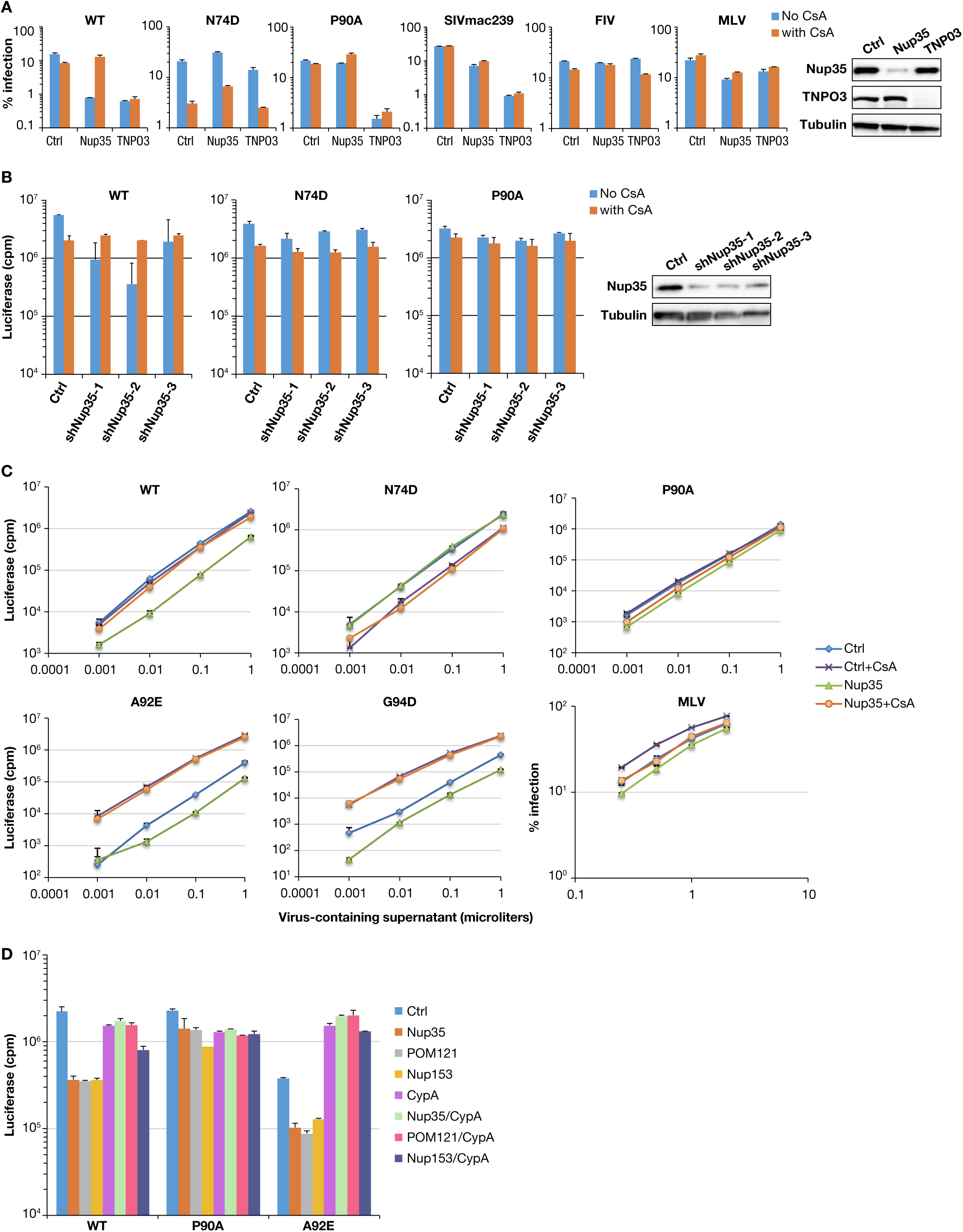
HIV-1 dependence on Nup35, Nup153, and POM121 is regulated by CypA. **a,** HIV-1, SIV, FIV, and MLV infection in the absence or presence of CsA in Nup35-depleted cells. HeLa cells were transfected with control siRNA or siRNAs against Nup35 for 48 h, then infected with VSV-G-pseudotyped viruses in the presence or absence of 2.5 μM CsA. After 48 h, infected GFP/RFP-positive cells were counted by FACS. Error bars show standard deviations of duplicates, representative of two independent experiments. Western blotting confirmed knockdown efficiency. **b**, CsA restores HIV-1 infection in Nup35-depleted MT4 cells. MT4 cells were transduced with a control shRNA or three different shRNAs against Nup35 then infected with VSV-G-pseudotyped WT or virus harboring various CA mutants. Firefly luciferase activity was measured at 48 h. Error bars show standard deviations of duplicates, representative of two independent experiments. Western blot analysis confirmed Nup35 depletion by three different shRNAs. **c**, CsA restores A92E and G94D HIV-1 infectivity in Nup35 knockdown cells. HeLa cells were transfected with control siRNA or siRNA against Nup35 for 48 h, then infected with increasing amounts of VSV-G-pseudotyped viruses in the presence or absence of 2.5 μM CsA, and firefly luciferase infectivity was measured at 48 h. Error bars show standard deviations of duplicates, representative of three independent experiments. **d**, CypA depletion by siRNA restores HIV-1 infectivity in Nup35 and POM121 knockdown HeLa cells. HeLa cells were either single-knocked down (Nup35, POM121, Nup153, or CypA) or double-knocked down (CypA in combination with Nup35, POM121 or Nup153) for 48 h. The cells were infected with VSV-G-pseudotyped pNL4-3-Luc-E-R+ or CA mutants (P90A or A92E) and firefly luciferase infectivity was measured at 48 h. MLV infection was also measured at 48 h in parallel samples to monitor cell toxicity and specificity of HIV-1 infection (Extended Data Fig. 5). Error bars show standard deviations of duplicates, representative of three independent experiments.

We sought to understand whether CypA affected WT HIV-1 dependence on Nup35 in the T cell line, MT4, which are highly permissive to transduction with shRNA-expressing lentiviral vectors. Cells were transduced with GFP-encoding shRNA vectors targeting Nup35 and then challenged with HIV-1 in the absence and presence of CsA (Fig. 2b). While knockdown of Nup35 specifically diminished WT HIV-1 infection in 2 of 3 shRNA lines, CsA treatment restored WT HIV-1 infectivity to levels observed in shRNA control cells treated with the drug. These data underline the pivotal role of CypA in determining the HIV-1 sensitivity to Nup35 depletion.

Given the interrelation between CypA binding and HIV-1 sensitivity to Nup35 depletion, we considered whether CsA-dependent viruses, A92E and G94D HIV-1, would remain sensitive to Nup35 depletion after CsA treatment. The CypA-dependent infection blocks that A92E and G94D HIV-1 exhibit in HeLa cells have been previously characterized as occurring at nuclear entry ^16^. In cells depleted of TNPO3, CsA treatment fails to fully restore the replication of these viruses indicating a qualitatively different infection block (data not shown). In comparison, A92E and G94D HIV-1 were also additionally sensitive to loss of Nup35 in cells (Fig. 2c). However in the presence of CsA, A92E and G94D HIV-1 infectivity in Nup35 knockdown cells rises to the level of the mutant viruses in control cells treated with CsA. Collectively, these data suggest the mechanisms underlying HIV-1 infection inhibition in Nup35 knockdown cells may be similar to the infection blocks encountered by A92E and G94D HIV-1 in normal cells in that CypA drives these viruses toward a nuclear entry pathway that they are unable to exploit.

## HIV-1 infection in POM121 and Nup153 knockdown cells is affected by CypA

The degree by which CypA controlled HIV-1 sensitivity to Nup35 depletion led us to investigate whether HIV-1 interactions with POM121, which emerged from the same screen, and Nup153 were CypA-dependent. Cells were either individually treated with siRNAs targeting Nup35, POM121, Nup153, and CypA, or doubly treated with siRNAs targeting Nup35, POM121, and Nup153 in combination with siRNA targeting CypA. Notably, depletion of CypA not only restored HIV-1 infection in Nup35 knockdown cells but similarly restored infection in POM121 knockdown cells and partly restored infection in Nup153 knockdown cells (Fig. 2d). A92E HIV-1 was also largely insensitive to the different Nup knockdowns in cells where CypA was also depleted. By contrast, MLV infection was largely unaffected in cells with single Nup knockdowns or Nup knockdowns in combination with CypA knockdowns (Extended Data Fig. 5).

## Nup35 knockdown impairs HIV-1 nuclear entry

To gain insight into the specific stage of HIV-1 infection at which Nup35 and CsA exert their effects, we performed time course experiments on Nup35-knockdown HeLa cells that were treated with CsA at various times after exposure to virus. Up to 12 hours post infection of Nup35-depleted cells, WT HIV-1 infectivity could be rescued by CsA treatment; however, CsA completely lost rescue ability at 24 hours post-infection (Fig. 3a). Similarly, CsA treatment also rescued the infectivity of CsA-dependent HIV-1 mutant A92E 12 hours post infection, but could not rescue infectivity at 24 hours post infection. P90A HIV-1 and MLV infection were not affected by CsA treatment at any time point, which is consistent with the inability of P90A HIV-1 and MLV to bind CypA.

**Figure 3.**
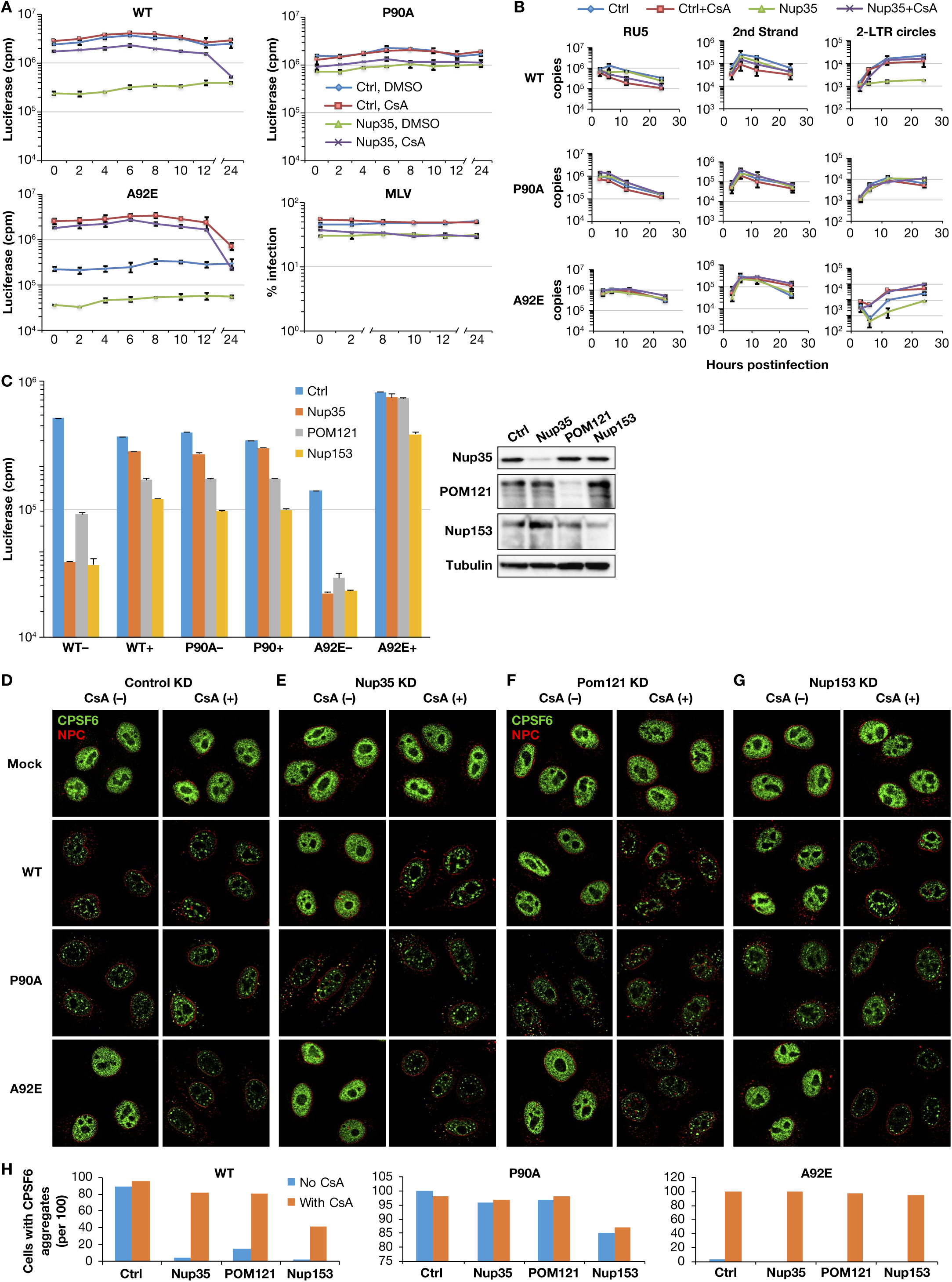
Nup35 knockdown impairs HIV-1 nuclear entry. **a,** CsA can restore HIV-1 infectivity in Nup35 knockdown cells 12 h after virus challenge. HeLa cells were transfected with control siRNA or siRNA against Nup35 for 48 h, then infected with VSV-G-pseudotyped viruses. CsA was added at every 2 h until 12 h post-infection and at 24 h post-infection. Firefly luciferase activity was measured at 48 h. Error bars show standard deviations of duplicates, representative of two independent experiments. **b**, HIV-1 reverse transcription is blocked in Nup35 knockdown cells but 2-LTR circle forms are not observed. HeLa cells were transfected with control or Nup35 siRNA for 48 h, then infected with VSV-G-pseudotyped, DNase-treated NL4-3-Luc-E-R+ or CA mutants (P90A or A92E) at an MOI of 1. The Cell DNA was extracted at 3, 6, 12, and 24 h after infection and used to detect early (RU5) RT, late (2nd strand) RT, and 2-LTR circles (right). Error bars show standard deviations of duplicates, representative of two independent experiments. The amount of DNA in each PCR assay was normalized by actin. HIV-1 infection was also measured at 48 h in parallel samples to monitor HIV-1 infectivity (Extended Data Fig. 6). **c,** CsA restores HIV-1 infection in Nup35, Nup153, and POM121 knockdown HeLa cells. HeLa cells were transfected with control siRNA or siRNAs against Nup35, POM121, or Nup153 for 48 h, then infected with infected with VSV-G-pseudotyped NL4-3-Luc-E-R+ or CA mutants (P90A or A92E) viruses in the presence or absence of 2.5 μM CsA. After 48 h, firefly luciferase infectivity was measured. Error bars show standard deviations of duplicates, representative of three independent experiments. Western blotting confirmed knockdown efficiency. - , no CsA; +, 2.5 uM CsA. **d-g**, CPSF6 distribution of Nup35, POM121, and Nup153 knockdown cells in the presence or absence of 2.5 μM CsA. Cells were infected with VSV-G-pseudotyped NL4-3-Luc-E-R+ or CA mutants (P90A or A92E) at an MOI of 50. After 12 h, cells were fixed and stained with antibodies against CPSF6 and NPC. **h**, The number of cells with or without CPSF6 aggregates were counted from (**d-g**).

Given the early reversibility of the infection block and the alteration of the nuclear pore complex by the various Nup knockdowns, we evaluated HIV-1 reverse transcription in cells depleted of Nup35 in the absence or presence of CsA (Extended Data Fig. 6 and Fig. 3b). We observed no difference in the accumulation of early or late reverse transcription products in control cells vs. Nup35-knockdown cells (Fig. 3b). Notably, both WT and A92E HIV-1 exhibited significantly reduced levels of 2-LTR circle junction forms of viral DNA (vDNA) unless treated with CsA. Although 2-LTR circular vDNA represents an abortive, noninfectious path, they form in the nucleus in the presence of DNA Ligase IV and thus can be used to assess viral nuclear entry^17^. These data indicate that the Nup35 knockdown does not impair reverse transcription but prevents HIV-1 nuclear entry in the presence of CypA.

We next tested whether CypA also regulated HIV-1 nuclear entry in cells depleted of POM121 or Nup153 using an assay based on CPSF6 distribution in HIV-1 infected cells. These cells were depleted of Nup35, POM121, or Nup153 in the absence of presence of CsA. As before, WT and A92E HIV-1 were sensitive to depletion of these Nups, and this infection reduction could be eliminated through CsA treatment (Fig. 3c). By contrast, MLV infection was not affected by CsA in Nup35, POM121, or Nup153 depleted cells (data not shown). Because CA interacts with CPSF6, soon after infection, it can induce CPSF6 precipitates, especially in the nucleus where CPSF6 is abundant (Fig. 3d). In Nup35-knockdown cells, WT HIV-1 infection no longer causes CPSF6 precipitation in the nucleus (Fig. 3e). Conversely, CsA treatment recovered CPSF6 aggregation following WT HIV-1 infection. As with the reverse-transcription staging data, these results indicate that restoration of HIV-1 infection by CsA is linked to nuclear entry. Similar results were obtained with A92E HIV-1. Notably, this virus did not induce nuclear aggregation of CPSF6 in control cells unless in the presence of CsA. As expected, CsA treatment did not alter CPSF6 distribution in P90A HIV-1 infected control cells, such that CPSF6 aggregates were observed in Nup35-knockdown cells in the absence of CsA. The phenotypes of WT HIV-1, P90A HIV-1, and A92E HIV-1 were consistent in cells depleted of either POM121 (Fig. 3f) or Nup153 (Fig. 3g). These results summarized in Fig. 3h indicate a pivotal role of CypA in regulating HIV-1 nuclear entry in Nup35, Nup153, and POM121 knockdown cells.

## The choreography of HIV-1 interaction with host factors governing nuclear entry reveals a regulatory switch

Given the pivotal role that soluble factor binding appeared to exert on transport of HIV-1 through the NPC, we considered whether differences in the subcellular localization of CPSF6 and CypA could predict their impact on this process. Although present throughout the cell, CypA is enriched in cytoplasmic compartments, especially in proximity to the nuclear membrane in both HeLa cells and T cell lines (Fig. 4a and Extended Data Fig. 7a-c). This distribution appears relatively stable and is not grossly affected by Nup35-depletion (Extended Data Fig. 7d). CPSF6, consistent with its role in pre-mRNA processing, is predominantly nuclear and distributed in a pattern quite distinct from CypA and is also unaffected by Nup35 depletion (Extended Data Fig. 7e).

**Figure. 4.**
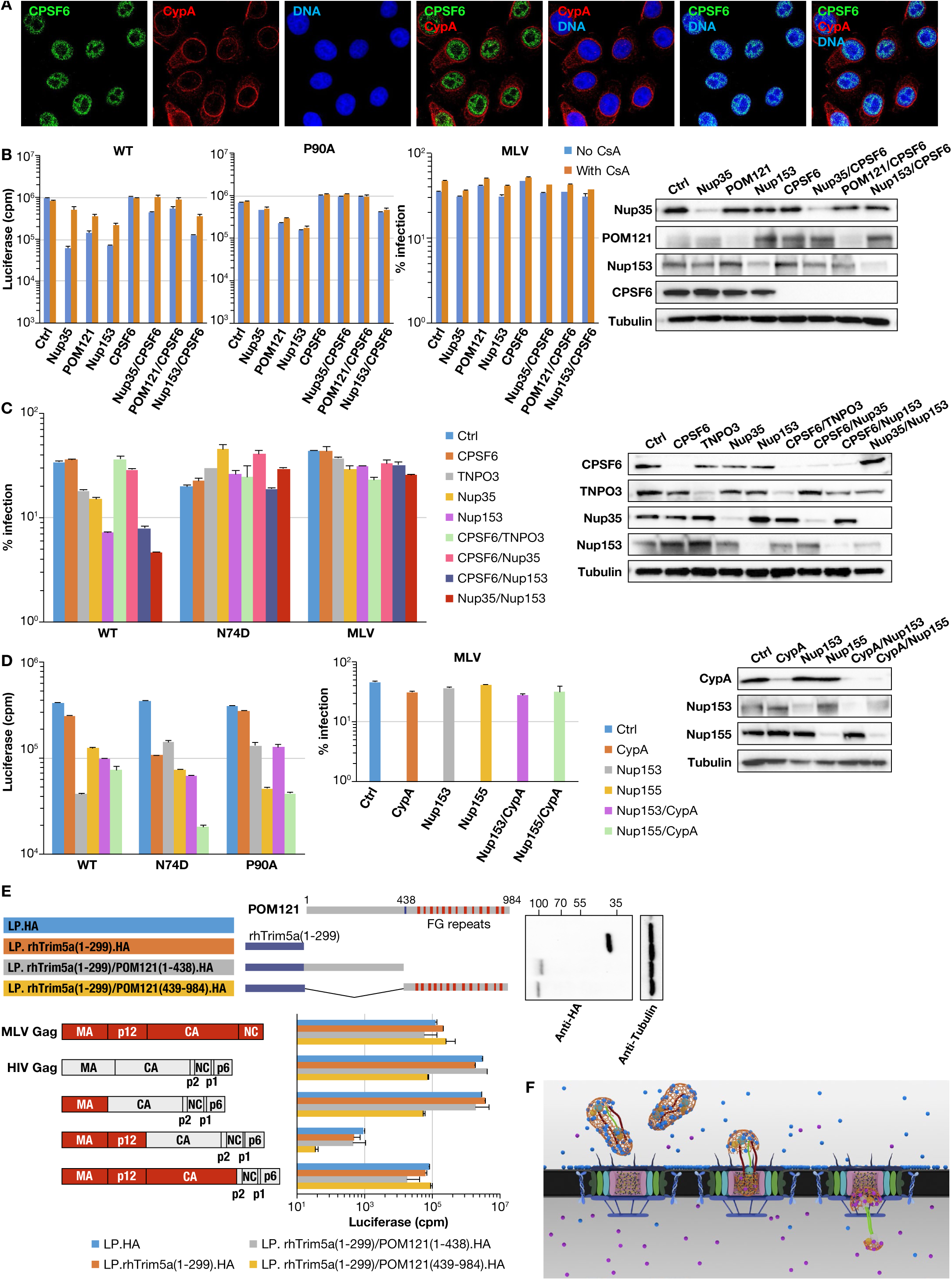
Soluble factors regulate nucleoporin dependence. **a,** CypA enrichment at the nuclear periphery. Localization of CPSF6 (green) and CypA (red) in HeLa cells in a Z-section image from deconvolution microscopy of immunostained HeLa cells. **b**, Partial restoration of HIV-1 infection in Nup35 and POM121 knockdown cells after CPSF6 depletion. HeLa cells were either single-knocked down (Nup35, POM121, Nup153, or CPSF6) or double-knocked down (CPSF6 with Nup35, POM121 or Nup153) for 48 h. The cells were infected with VSV-G-pseudotyped NL4-3-Luc-E-R+ or CA mutants (N74D or P90A) viruses in the presence or absence of 2.5 μM CsA, and firefly luciferase infectivity and percentage of RFP-positive cells (MLV infection) were measured at 48 h. Error bars show standard deviations of duplicates, representative of three independent experiments. Western blotting confirmed knockdown efficiency. **c**, Distinct contributions of Nup35 and Nup153 to HIV-1 infection. HeLa cells were either single-knocked down (CPSF6, TNPO3, Nup35, or Nup153) or double-knocked down (CPSF6 with TNPO3, Nup35, or Nup153; Nup35 with Nup153) for 48 h. The cells were infected with VSV-G-pseudotyped HIV-RFP or CA mutant (N74D) viruses, and percentage of RFP-positive cells were measured at 48 h. Error bars show standard deviations of duplicates, representative of three independent experiments. Western blotting confirmed knockdown efficiency. **d**, CypA determines HIV-1 dependency on Nup153 versus Nup155. HeLa cells were either single-knocked down (CypA, Nup153, or Nup155) or double-knocked down (CypA with Nup153 or Nup155) for 48 h. The cells were infected with VSV-G-pseudotyped pNL4-3-Luc-E-R+ or CA mutants (N74D or P90A) viruses, and firefly luciferase infectivity and percentage of RFP-positive cells (MLV infection) were measured at 48 h. Error bars show standard deviations of duplicates, representative of three independent experiments. Western blotting confirmed knockdown efficiency. **e**, TRIM5-POM121 inhibition of HIV-1 infection is CA-dependent. Top left, schematic representation of the rhesus TRIM5 RBCC (RING, B-box2, and coiled coil) domains (1-299) with the POM121 N-terminal fragments (1-438) or C-terminal fragments (439-984). Top right, western blotting analysis of lysates from HeLa cells expressing TRIM fusion proteins. Bottom left, schematic representation of the chimeric HIV/MLV virus. Bottom right, HeLa cells stably transduced with HA-tagged rhTrim5a/POM121 fusion constructs were infected with VSV-G-pseudotyped chimeric HIV/MLV virus, and firefly luciferase infectivity was measured at 48 h. Error bars show standard deviations of duplicates, representative of three independent experiments. Western blot analysis of HA-tagged rhTrim5a/POM121 fusion and control constructs in stable HeLa cells (Extended Data Fig. 8). **f**, CypA determines the HIV-1 nuclear import pathway. Premature CPSF6 interaction with HIV-1 impairs CA interaction with FG-Nups at the NPC. CypA use by HIV-1, however, prevents CPSF6 access to the N74 pocket in CA until the virus docks at the NPC. Subsequent CPSF6 binding may enhance release from the NPC. We propose that the HIV-1 core, comprised of multimeric CA in association with the viral nucleic acid and enzymatic proteins, directly functions as a NTR and exploits successive FG interactions to achieve transfer through the NPC but these interactions are regulated in time and space by CypA and CPSF6.

Indeed, in cells depleted of Nup35, POM121, or Nup153, only a partial restoration of WT HIV-1 infection is observed after coordinate knockdown of CPSF6 (Fig. 4b), especially in Nup35 and POM121 knockdown cells. HIV-1 infection in these cells can be fully restored after CsA treatment. In comparison, P90A HIV-1 does not interact with CypA, and the infectivity of this virus is slightly diminished in POM121 and Nup153 knockdown cells (Fig. 1a, Fig. 3c, and Fig. 4b). Whereas CsA treatment, predictably, does little to affect P90A infection in the POM121 or Nup153 knockdown cells, CPSF6 depletion potently enhances infection when in conjunction with either knockdown (Fig. 4b).

Although WT HIV-1 infection in Nup35 knockdown cells was elevated after CPSF6 knockdown, infection in Nup153 knockdown cells remained impaired even after CPSF6 knockdown, suggesting distinct contributions of both Nups to HIV-1 nuclear entry. These observations were confirmed and extended. While the effect of TNPO3 knockdown on HIV-1 infection could be significantly ameliorated by CPSF6 depletion, and the effect of Nup35 knockdown on HIV-1 infection could be partially offset by CPSF6 depletion, the reduction in HIV-1 infection by Nup153 knockdown was unaffected by CPSF6 depletion (Fig. 4c). Notably, coordinate depletion of Nup35 and Nup153 potently impaired WT but not N74D HIV-1 infection.

We next asked whether CypA was also a key regulator of Nup155 use. Nup155 is a non-FG Nup in the Nup93-subcomplex that we previously noted N74D HIV-1 to be more dependent for infection than WT HIV-1^2^. WT HIV-1 infection is diminished in cells depleted of Nup153 but retains greater infectivity in Nup155 knockdown cells (Fig. 4d). Two distinct effects on infection are observed when CypA is depleted in Nup153 and Nup155 knockdown cells. WT HIV-1 infection is increased in the Nup153 knockdown cells as previously noted, and it is decreased in the Nup155 knockdown cells (Fig. 4d). The infectivity of WT HIV-1 in the CypA plus Nup155 double knockdown cells is comparable to N74D and P90A HIV-1 infection of the Nup155 cells. In the absence of CypA, it becomes more dependent on Nup155. These data suggest the use of a secondary nuclear pathway for HIV-1 when insertion into the Nup153 pathway is not possible.

Although our data suggest that the CA-dependence for HIV-1 use of Nup35 and POM121 is linked to soluble host factor binding, it remains possible that these proteins also make direct contact with HIV-1 despite their position within the NPC. We thus sought to examine whether fragments of Nup35 or POM121 could interfere with HIV-1 in a CA-dependent manner. This approach has been leveraged in mapping CA-interaction domains in CPSF6 and Nup153, specifically through fusion to the N-terminal RING, B-box 2, and coiled-coil (RBCC) domains of rhesus TRIM5 alpha, substituting the B30.2 (SPRY) domain with a region of interest ^10,18^. Attempts to make rhTRIM5-Nup35 fusions that were detectable in cell lines were unsuccessful. By contrast, rhTRIM5 fusions to the N-terminus and C-terminus of POM121 were stable (Fig. 4e). Due to the potency of rhTRIM5-mediated restriction and the sensitivity of WT, N74D, and P90A HIV-1 to POM121 depletion (Fig. 1a), we first tested whether HIV-1 and MLV were differentially susceptible to infection in cells expressing the TRIM5-POM121 fusion proteins (Fig. 4e). Whereas MLV displayed no apparent sensitivity to the different TRIM5-POM121 proteins, HIV-1 (LAI Gag) was specifically inhibited by a TRIM5 fusion to the FG-rich, C-terminal domain of POM121. Moreover, testing with HIV-1/MLV Gag chimeric viruses revealed that HIV-1 CA was required for sensitivity to this protein. We extended this analysis to the CA-mutant HIV-1 (NL4-3 Gag) isolates (Extended Data Fig. 8). Although HIV-1 CA governs susceptibility to TRIM5-POM121-Cterm, the N74D and P90Amutations were insufficient to escape restriction by the fusion protein.

## Discussion

Here we demonstrate the dependence of HIV-1 on Nup35 and POM121 for nuclear entry. As has been observed for Nup153 and Nup358, HIV-1 CA determines sensitivity to loss of Nup35 or POM121. Extending prior work, soluble CA-interacting factors also regulate the necessity for the nucleoporins. Unlike smaller cargo that interacts with cellular NTRs to achieve nuclear entry, we propose that the HIV-1 core, comprised of multimeric CA in association with the viral nucleic acid and enzymatic proteins, directly functions as a macromolecular NTR and negotiates multiple nucleoporin interactions to achieve transfer through the NPC, through a pathway determined by soluble host factor binding in the cytoplasm. Like cellular NTRs, CA selectively interacts with FG dipeptides ^10,12^. In contrast to cellular NTRs, HIV-1 can enter the nucleus of metabolically inert cells without being dependent on Ran-GTP regulation. Given that the host cell cytoplasm contains sensors to detect and interfere with microbial pathogens, slipping into the nucleus quickly is advantageous to the virus.

Although only 24 kilodaltons in size as a monomer, the hundreds of CA molecules present in a hexameric lattice composing the cytoplasmic HIV-1 core likely enable a distribution of protein-protein interactions with host factors based on local concentrations and relative binding affinities. Human host factors present in the cytoplasm known to interact with CA include CPSF6, CypA, and MxB. MxB is thought to be enriched proximal to NPCs after interferon activation of cells ^19^. By contrast, CPSF6 and CypA are present at high steady levels, with CPSF6 predominantly enriched in the nucleus and CypA abundant in the cytoplasm, especially in proximity to the nuclear membrane. HIV-1 has likely evolved to exploit this concentration and distribution difference between CPSF6 and CypA in different compartments to enhance access of NPCs, to ensure binding surfaces on CA were available for interaction with specific Nups, and to utilize CPSF6 in the nucleus to access gene-rich regions of chromatin. Prior studies have indicated that it is detrimental to the virus to engage these factors out of order. Enrichment of CPSF6 in the cell cytoplasm blocks HIV-1 nuclear entry presumably by preventing Nup interactions with the virus ^2,20^.

The role of CypA in HIV-1 infection has been a long-standing puzzle. It was one of the first host factors identified to interact with the Gag protein of an animal retrovirus ^21^. Although it had been mapped to exert its effect on early replication steps prior to nuclear entry ^22,23^, the precise mechanism has been unclear. Studies with Nup153 and Nup358 suggested that CypA could affect subsequent interactions with these Nups ^5,24^, but to a limited extent. It is possible that the essential role that these Nups play in the transport of all cargo limits analysis after they are depleted from cells. By contrast, the effect of CypA on HIV-1 dependence of Nup35 or POM121 was dramatic. While HIV-1 was blocked for infection in Nup35 or POM121 knockdown cells, after treatment with CsA, HIV-1 enters the nucleus and infection is restored to levels of control cells. CypA-binding thus directed HIV-1 to a nonproductive nuclear import pathway in the knockdown cells, possibly one that starts with Nup358 interaction. Thus, CypA might not only block premature CPSF6 interaction before docking at the pore, its association with HIV-1 might facilitate a transfer of the core to Nup358, initiating a specific path of passage through the NPC.

At this point, it is unknown whether Nup35 and POM121 make direct contact with HIV-1. Nup35 is a member of the Nup93 subcomplex which encircles the central channel Nup62 subcomplex comprised entirely of FG-Nups. Nup35 itself has three FG dipeptides, but it is unknown if these are accessible to cargo. Given the demonstrated interaction of FG dipeptides in CPSF6 and Nup153 with the N74 pocket of HIV-1 CA, it is tempting to speculate that this interface of HIV-1 negotiates a multitude of interactions by the core during passage through the NPC. To a large extent, Nup35 and POM121 knockdowns phenocopied Nup153 knockdowns with regards to CA-dependent effects on HIV-1 infection. Similar to Nup153 and CPSF6, TRIM5 fusions to the FG-rich portion of POM121 inhibit HIV-1 infection. Recently, another group has also shown that a fragment of highly related POM121C containing numerous FG-motifs was sufficient to impair HIV-1 infection ^25^. While they did not investigate whether POM121/POM121C could serve as a co-factor through knockdown/knockout experiments, they argue that overexpression of truncated POM121C binds to a subunit of a cellular NTR, Karyopherin subunit beta-1 (KPNB1), which possibly underlies a block to HIV-1 infection. KPNB1 forms a complex with Importin-7 to mediate transport of cargo to the nucleus^26^. KPNB1 interaction with POM121C is not unexpected given that NTRs interact with FG-Nups. More recent studies have not supported a role for Importin-7 in HIV-1 infection ^27^. Consistent with this finding, we also did not observe removal of either KPNB1 or Importin-7 to effect HIV-1 infection (Extended Data Fig. 9). Instead POM121 interaction with CA appears to support HIV-1 infection.

Notably, POM121 is known to interact with Sun1 ^28^. Sun1 and Sun2 combine to form a linker of nucleoskeleton and cytoskeleton (LINC) complexes which span the nuclear envelope ^29^. Knockdown or knockout of Sun2 reduces HIV-1 infection in human primary CD4+ T cells ^30^ and THP-1 cells ^31^, respectively. The CypA-dependence of Sun2 is unsettled. One study has shown that Sun2 promotes CypA-dependent steps of HIV-1 replication in bone marrow-derived dendritic cells from *sun2-/-* mice^30^, but other two studies have shown that there is no correlation between Sun2 and CypA in human primary CD4+ T cells ^32^ and THP-1 cells ^31^. These differences could be due to cell context. Sun1 and Sun2 form an inner nuclear membrane (INM) complex proximal to NPCs. It will be important to determine the contribution of INM proteins with nucleoporins in influencing HIV-1 nuclear entry pathway. How Sun2 interacts with HIV-1 is a subject of investigation.

Elucidating the precise choreography of HIV-1 interactions at the nuclear membrane and during passage through the NPC is critical to understanding the successive contributions of each of these factors. As illustrated in Figure 4c, the Nup35 and Nup153 blocks to HIV-1 infection were additive in double-knockdown cells. One interpretation of these data is that the blocks are distinct and Nup35 loss does not entirely prevent Nup153 use by HIV-1. It could be argued that the removal of both factors was incomplete, so the effect of the double knockdown was increased potency in inhibiting the same pathway. However, the genetics of both blocks are distinct. While there may be an overlapping need for function by both Nup35 and Nup153 for optimal HIV-1 nuclear entry and infection, CPSF6, potentially interacting with the virus within the NPC, creates a more stringent requirement on Nup35 (Fig. 4c) indicating functional differences between successive steps during nuclear import.

Indeed our prior work pointed to the existence of distinct routes of nuclear entry available to HIV-1 ^2^, which split along an axis of WT vs N74D HIV-1 CA exhibiting greater dependence on Nup153 vs Nup155, respectively. We find here that this switch in pathway use is governed in part by CypA. N74D HIV-1 exhibits great sensitivity to the loss of CypA with an infection block in the cytoplasm ^15^. When Nup155 loss is coupled with CypA depletion in experiments presented here, a greater than 50-fold decrease in N74D HIV-1 infectivity is observed. Moreover, WT HIV-1 also becomes more dependent on the Nup155-import pathway in CypA-depleted cells. For WT HIV-1, a loss of CypA binding diminishes access to the Nup153-import pathway. For N74D HIV-1, the N74D mutation of CA precludes Nup153 utilization.

A competition between CypA and CPSF6 for binding to HIV-1 CA likely exists in the cell cytoplasm given the overlap of binding sites. CypA is one of the most abundant proteins in the cell. With the elevated concentration of CypA proximal the nuclear membrane, HIV-1 may have evolved to bind it to prevent premature interaction with CPSF6 (Fig. 4f). CPSF6 aggregates are in fact observed in the cytoplasm of cells infected with P90A HIV-1 (Fig. 3f-g). The affinity of full-length CPSF6 interaction with CA hexamers is currently unknown, although a peptide of the CA-binding site of CPSF6 has an estimated affinity (K_D_) of 50 micromolar ^4^. CypA binding to the CA lattice at an affinity (K_D_) of approximately 12 micromolar ^33^. Thus it is unlikely that CPSF6 will exhibit preferential binding to cores in areas of the cell where CypA is at higher concentration.

While both HIV-1 and SIVsm/mac/mne viruses retain CPSF6 binding interfaces, HIV-1 distinctly acquired a CypA interaction site in proximity to the CPSF6 binding interface. This potentially reflects differences in the subcellular localization of CPSF6 in human vs some nonhuman primate cells. If human cells have elevated CPSF6 levels outside the nucleus, this could impair HIV-1 interaction with the NPC. As we observe here (e.g. Fig. 2a,b) and has been noted in past studies ^13,14^, CsA treatment of some human cell targets results in diminished WT HIV-1 infection. Despite the potential danger of cytoplasmic CPSF6 to the virus, CPSF6 has a positive effect on Nup35 and POM121 contribution to HIV-1 infection (Fig. 4b,c). How to reconcile both CypA and CPSF6 effects on HIV-1 infection when there might be competition for binding HIV-1 CA? We suggest a transition occurs after HIV-1 interaction with Nup358 at the cytoplasmic face of the NPC, previously hypothesized by Schaller et al ^5^. Some CypA is likely displaced from the core by the Nup358-CypA related domain, which provides CPSF6 and probably a subset of FG-Nups to subsequently interact with CA hexamers. This would explain why the N74D HIV-1 infection pathway is different, resembling the entry route of other lentiviruses, such as FIV ^2,24^. It fails to interact with CPSF6 within the NPC and uses distinct, currently unidentified, surfaces on CA to negotiate potential nucleoporin interactions.

Despite WT and N74D HIV-1 exhibiting differences in their use of the NPC, there are common themes. Removal of Nup62 subcomplex members (Nup54, Nup58, and Nup62), which are FG-Nups that form a meshwork in the central channel of the pore restricting the flow of cargo larger than 5 nm in diameter, generally elevated both WT and N74D HIV-1 infection. While loss of Nup54 and Nup58 elevated infection of WT HIV-1, N74D HIV-1, and P90A HIV-1, removal of Nup62 had a larger positive effect on WT HIV-1 infection. Knockdown of Nup214, a FG-Nup that also is thought to regulate the flow of cargo, also had a greater positive effect on WT HIV-1 infection. Collectively, WT HIV-1 infection appears to be more broadly regulated by FG-Nups. Although we did not directly investigate the role of CypA in facilitating Nup62 and Nup214 interaction, P90A HIV-1 infection was 2-3-fold lower than WT HIV-1 infection under knockdown conditions.

The ability of CypA/CPSF6 to regulate HIV-1 interactions with the NPC underscores the vulnerability of the virus at this crucial replication step to MxB which also targets HIV-1 CA ^34–36^. Both N74D^34,35^ and P90A^34^ HIV-1 show increased resistance to MxB antiviral function. Similarly, other CypA and CPSF6 CA-binding mutants G89V and N57S, respectively, also exhibit MxB resistance ^35^. MxB restriction has been associated with CypA function^36^, and it is proposed that the block occurs at the nuclear entry ^34,35^. The potential interplay of MxB with CypA or CPSF6, and whether it dysregulates the temporal or spatial sequence of their interactions with CA, has not been examined at this point.

CA is intrinsic to critical replication steps in the cytoplasm, at the nuclear membrane, and within the nucleus. Many of these actions are likely dependent on retention of a portion of the core, especially given the N74 interface is accessed by factors when CA is in a hexameric configuration. The coordination of different successive interactions among a limited subset of interfaces helps explain why that this choreography is profoundly sensitive to disruption by small molecule inhibitors targeting CA, such as PF-74 and BI-2^37,38^.

Taken together, this study supports a model of HIV-1 CA co-opting CypA as a key factor in preventing premature interactions in the cytoplasm by CPSF6 and presumably newly synthesized, soluble nucleoporins until the virus reaches the nuclear membrane. Subsequently, the CA lattice functions as a NTR which binds to different FG motifs present in FG-Nups or CPSF6 in the nuclear channel enabling HIV-1 transport into the nucleus. The transposon Tf1 is similarly hypothesized to use an assembled Gag particle as a multimeric NTR, particularly in interacting with the FG-rich yeast Nup124p during nuclear entry ^39^. With regards to animal cell biology, as our and previous studies indicate, HIV-1 provides an attractive model to interrogate the mechanism of nuclear entry by large macromolecular entities.

## METHODS

### Plasmids

pNL4-3-Luc-E-R+ (HIV-1 vpr-positive, env-deleted, encoding firefly luciferase in place of nef) or pHIV-RFP (HIV-1 vpr-negative, env-deleted, encoding RFP in place of nef) were used in transfections with pL-VSV-G to generate VSV-G-pseudotyped HIV-1 vectors, and the MLV-based retroviral vector pMX-RFP was used to generate Moloney-based virus in co-transfections with pJK3, pL-VSV-G, and pCMV-Tat. Vectors for FIV ^40^ and SIV_MAC239_ ^41^ have been described previously.

cDNA encoding human Nup35 was amplified from MGC Human Nup35 sequence-verified cDNA (Accession number BC047029, GE Healthcare, Dharmacon), using primers that add an amino terminal HA tag, and inserted into pLPCX-MCS between the EcoRI and NotI sites. All coding sequences were confirmed by DNA sequencing.

### Cells and culture conditions

HEK293T and HeLa cell lines were maintained in DMEM supplemented with 10% FBS. H9, Jurkat, and MT4 cells were grown in RPMI 1640 plus 10% FBS. GHOST cells were maintained in DMEM supplemented with 10% FBS, 500 ug/ml G418, 100 ug/ml hygromycin, and 1 ug/ml puromycin.

To generate stable Nup35-expressing cell lines, HeLa cells were transduced with either wild-type or mutant HA-tagged Nup35, a pLPCX-HA-Nup35 (cds) vector containing only the amino acid coding sequence of the human Nup35 cDNA, which excludes the 3’-UTR, and the cells were selected with 1 μg/ml puromycin (Millipore).

### siRNA screen

To identify Nups requirements for HIV-1 infection, 32 human Nups were systematically depleted using two different commercially available siRNA pools (Dharmacon; Sigma). siRNAs were transfected into the HeLa cells at a 50 nM final concentration using RNAiMAX (ThermoFisher Scientific), according to the manufacturer’s instructions. After 2 days, the cells were re-seeded into 24-well plates and infected with VSV-G-pseudotyped RFP reporter viruses at an MOI of 0.5 in the presence of 5 ug/ml polybrene. After an additional 48 h incubation, cells were trypsinized and the percentage of RFP-positive cells was scored by flow cytometry (FACSCalibur, BD Biosciences). As a positive control, siRNA SMARTpool against TNPO3 (Dharmacon) was present on each plate. To exclude false positives (e.g., due to toxicity), WT HIV-1, N74D HIV-1, and P90A HIV-1 infections were performed in parallel to Moloney murine leukemia virus (MLV) infection on the Nup-depleted cells. The first round of the screens was performed using siRNA from Sigma. Those genes that showed a phenotype in the first round were confirmed by Dharmacon siRNA. Therefore, two-thirds of the genes were screened using both Dharmacon on-target siRNA SMARTpool and Sigma MISSION esiRNA. The screening was repeated in at least three independent experiments.

### Viral production

To produce HIV-1 particles, pNL4-3-Luc-E-R+ or HIV-RFP (wild-type and capsid mutants: N57A, Q63/67A, K70A, N74D, P90A, A92E, G94D, and T107A) was co-transfected with a VSV-G expression vector at a ratio of 3:1 using Hilymax (Dojindo Molecular Technologies). The medium was replaced after overnight incubation and viral supernatants were collected at 48 h post-transfection. Viral supernatants were filtered and their infectivity was determined by using HeLa or GHOST target cells.

MLV stocks were obtained by co-transfection of pLPCX, pJK3, pL-VSV-G, and pCMV-Tat at a ratio of 4:2:1:0.3, respectively, with Hilymax. The medium was replaced one day after overnight incubation and viruses were harvested at 48 h after transfection, passed through a 0.45-μm filter, and used directly to transduce target cells.

### Viral infection

For single cycle infectivity assays, HeLa cells were plated in 24-well plates at 5×10^4^ cells per well, and infected with single or serial-dilutions of VSV-G-pseudotyped HIV-1, SIVmac_239_, FIV, or MLV in the presence of 5 ug/ml polybrene. In some experiments, 2.5 μM cyclosporine A (Bedford laboratories) was added to the culture media at the time of infection. At 48 h post-infection, cells were washed, and the infection was analyzed by examining the percentage of RFP or GFP expressing cells using flow cytometry (FACSCalibur, BD Biosciences), or by measuring firefly luciferase activity (Promega).

For microscopy experiments, 2.5×10^4^ HeLa cells were grown overnight on glass slides in a 24-well plate. The next day, cells were infected with WT, P90A, or A92E pNL4-3-Luc-E-R+ at an MOI of 50. At 12 h post-infection, cells were washed three times with PBS, fixed, permeabilized, and stained with antibodies.

For analysis of HIV-1 reverse transcription products, HeLa cells were seeded at 2×10^5^ cells per well in 6-well plates and infected with VSV-G-pseudotyped wild-type HIV-1 or capsid mutant virus (MOI of 1) in the presence or absence of 2.5 μM cyclosporine A. For the negative control, a reverse transcription inhibitor (EFV 150 nM) was added at the time of infection. The phenotype infection assay was performed in parallel in 24-well plates.

### RNA interference

Three ON-TARGETplus siRNAs directed against human Nup35 were purchased from Dharmacon: siRNA 1, 5’-CUGCUGGUUCCUUCGGUUA-3’; siRNA 2, 5’-AGAUAAAAGUGGCGCUCCA-3’; siRNA 3, 5’-AGUUAUUUCUACC GACACA-3’. HeLa cells were plated at 1.5×10^5^ cells per well in 6-well plates and transfected the next day with a final concentration of 40 nM siRNA targeting Nup35 or non-targeting control siRNA (siCONTROL non-targeting siRNA, Dharmacon), using RNAiMAX (ThermoFisher Scientific) according to the manufacturer’s instructions. To generate stable Nup35 knockdown cells, a lentiviral vector and the following GIPZ Lentiviral (GE Healthcare, Dharmacon) shRNAs against hNup35 were used in this study:

GIPZ Lentiviral Human Nup35 shRNA:

V3LHS_364366, TGTCTGTCAGAAATAACCT
V3LHS_380787, TATGAGCTGGTACAACTGG
V3LHS_380788, TGGTTCAGATCCTAACGCG

Lentiviral vector stocks were produced by co-transfection of 293T cells with packaging plasmid ΔR8.2, MISSION shRNA or GIPZ Lentiviral shRNA, and pL-VSV-G at a ratio of 1:1:0.5, the culture medium was replaced at 12 h, and viral supernatants were collected and filtered at 48 h. HeLa cells were transduced with the filtered supernatant and selected in 2 ug/ml puromycin.

### CRISPR-Cas9 knockout

To generate a CRISPR-Cas9 single guide RNA (sgRNA) expression vector, oligonucleotides were annealed and inserted into the pX330 (Addgene plasmid #42230). The following sequences targeting coding regions at the 5’ end of the gene were used: 5’-GAAGGGCCACTAATTGATCG-3’ (Nup35, exon3) and 5’-GGACGCGGCGTTGCCACCAG-3’ (Nup153, exon1). Non-targeting sgRNA was used as control, 5’-CGCTTCCGCGGCCCGTTCAA-3’. HeLa knockout cells were generated by transiently transfecting pX330 into the cells. Single clones were screened by western blotting and phenotype assay and the candidate clones were verified by sequencing of genomic DNA using the following primers: 5’-cgGAATTCACATCTCCAAAGCCAGGAGTTA-3’ and 5’- taAAGCTTTGTATTTTATGTGTGGCCCAAG-3’ (Nup35 target exon3), 5’- cgGAATTCCTCTAAGGCCTCCGCCTCT-3’ and 5’- taAAGCTTGTCCCATACCTGATGCTGTTGT-3’ (Nup153 target exon1).

### RNAi-resistant mutant generation and phenotype rescue

To generate RNAi-resistant variants of genes, the exogenous ORF transcript lacking the 3’-UTR sequence targeted by the siRNA was inserted into LPCX vector. HeLa cells were then transduced with the filtered supernatant and the cells were selected with 1 μg/ml puromycin (Millipore). For the rescue assay, puromycin-selected cells were plated at 1.5×10^5^ cells per well in 6-well plates and transfected the next day with siRNA (40 nM) against human Nup35 3’-UTR region or control siRNA. Two days post-transfection, cells were re-seeded and infected with the HIV-1 or MLV and analyzed by FACS 2 days post-infection.

### Quantification of HIV-1 reverse transcription products by real-time PCR

HeLa cells were transfected with siRNAs as described above. Two days later, cells were seeded at 2×10^5^ cells per well in 6-well plates and infected with VSV-G-pseudotyped HIV-1, or capsid mutant virus P90A HIV-1, or A92E HIV-1. The virus was pretreated with 20 U ml^-1^ RNase-free DNase I (Roche) for 1 h at 37°C. At 2 h post-infection, the cells were washed once with PBS and fresh complete medium was added, cells were then collected at 3, 6, 12 and 24 h after infection. Total DNA was extracted using the QIAamp DNA Blood Mini Kit (Qiagen) and used for real-time PCR to specifically quantify HIV early reverse transcription (RT) products, late RT products, and 2-LTR circle forms. The primer-probe sets and conditions were used as previously described ^42^. To normalize the amount of DNA in each PCR assay, the following primer set was used to quantify the copy number of the cellular gene actin: forward, 5’- TCACCCACACTGTGCCCATCTA CGA -3’ and reverse 5’- CAGCGGAACCGCTCATTGCCAATGG -3’. qPCR assays were performed using either Platinum qPCR SuperMix-UDG (Invitrogen) for detecting HIV-1 reverse transcription or iQ SYBR Green Supermix (Bio-Rad) for detecting actin.

### Western blotting

Whole-cell extracts were prepared by lysing cells in RIPA buffer (Sigma-Aldrich), equivalent protein content boiled in SDS sample buffer, resolved by Criterion Tris-HCl Precast Gels (Bio-Rad), and blotted onto PVDF Blotting Membranes (GE Healthcare). Membranes were probed with primary antibodies specific for human cyclophilin A (Abcam), CPSF6 (Novusbio), Importin 7 (Novusbio), KPNB1 (Novusbio), Nup35/53 (Abcam, Bethyl, GeneTex, or Novusbio), Nup93 (Abcam), Nup153 (Abcam), Nup155 (Abcam), Nup188 (Novusbio), Nup205 (Novusbio), Nup358 (Abcam), POM121 (GeneTex), TNPO3 (MyBioSource), HA (Sigma), and tubulin (Sigma) followed by secondary HRP-conjugated anti-mouse (GE Healthcare) or anti-rabbit (GE Healthcare) antibodies and detected using a Chemidoc XRS+ system (Bio-Rad).

### Immunofluorescence

Cells were fixed with 4% paraformaldehyde (Boston Bioproducts) at room temperature (RT) for 10 min, permeabilized with 0.2% Triton X-100 in PBS for 10 min, and then blocked with 3% BSA for 30 min. Cells were incubated with primary antibodies at RT for 1 h, followed by Alexa-Fluor-conjugated secondary antibodies and Hoechst 33342 (Thermo Fisher Scientific) at RT for 30 min. Coverslips were mounted on glass slides with ProLong Gold antifade solutions (Molecular Probes). Images were taken in Z-stacks using the Deltavison deconvolve microscope (GE Healthcare) and deconvolved to remove out-of-focus light using the Softworks software (GE Healthcare). Primary and secondary antibodies used for immunofluorescence were mouse anti-Cyclophilin A (Abcam), rabbit anti-CPSF6 (Novusbio), mouse anti-Nup153 (Abcam), Rabbit anti-Nup155 (Abcam), mouse anti-Nuclear Pore Complex (MAb 414, BioLegend), rabbit anti-Lamin B1 (Abcam), goat anti-rabbit Alexa 488 (Molecular Probes), goat anti-mouse 488 (Molecular Probes), goat anti-mouse 546 (Molecular Probes), and goat anti-rabbit 555 (Molecular Probes).

## Acknowledgements

We thank Szu-Wei Huang and KyeongEun Lee for intellectual contributions and scientific discussions. We thank Eric Freed, Henry Levin, and Owen Pornillos for scientific suggestions, and Eric Freed for MT4 cells. We thank Joseph Meyer from Scientific Publications, Graphics and Media, Frederick National Laboratory for assistance with figure graphics. Dr. Xue was an NIH Intramural AIDS Research Fellowship and NCI Sallie Rosen Kaplan Award recipient for women scientists. This research was supported by the Intramural Research Program of the NIH, Frederick National Lab, Center for Cancer Research. This work was also supported by NIH grants 5R01AI111809, 5DP1DA034990, and 1R01AI117839, to J.L. S.G.S. acknowledges support by NIH grants P50 GM103368 and R01 AI120860. The content of this publication does not necessarily reflect the views or policies of the Department of Health and Human Services, nor does mention of trade names, commercial products, or organizations imply endorsement by the U.S. Government.

## Extended Data Legends

**Extended Data Figure 1: Depiction of the nuclear pore complex and HIV-1 CA hexamers. a**, Major human NPC subcomplexes. FG-Nups hypothesized to interact with nuclear pore cargo based on yeast or metazoan studies are emboldened. Image adapted from Ori *et al*^6^. **b and c**, Overlap of CypA and CPSF6 binding regions in the HIV-1 capsid. Surface representation of the structure of two adjacent hexamers from the cryo-EM model of capsid (CA) tubes (PDB ID: 3J34; yellow CA_NTDS_ and orange CA_CTDS_). (**b**) Binding site of a CPSF6 peptide (residues 313-327; magenta) based on crystal structures PDB ID: 4U0A and 4WYM. (**c**) Binding site of CypA based on cryo-EM model PDB ID: 5FJB; blue. The structures suggest that CypA bound to capsid may affect access of the full size CPSF6 to its binding site.

**Extended Data Figure 2: Effect of nucleoporin knockdown on MLV infection.** HeLa cells were transfected with a control siRNA or target siRNA (smart pool) for 48 h, then infected with m.o.i of 0.5 of VSV-G-pseudotyped MLV. After 48 h, infected RFP-positive cells were counted by FACS. This analysis was performed in parallel the experiments shown in Fig. 1a. Percent relative infection is mean ± s.d., from three independent experiments. MLV, murine leukaemia virus.

**Extended Data Figure 3: Nup35 knockdown does not affect levels of other nucleoporins.** Nup35 knockdown does not affect levels of other proteins in the Nup93 subcomplex. Western blot analysis with anti-Nup35 (**a**) or antibodies against Nup93 subcomplex proteins (**b**) using lysates from Nup35-knockdown HeLa cells. **c,** Nup153 or Nup358 levels are unchanged in Nup35 knockdown cells. Western blot analysis showing reactivity to anti-Nup35 or antibodies against TNPO3, Nup153, and Nup358 using Nup35 or TNPO3 knockdown HeLa cell lysates.

**Extended Data Figure 4: Generation of knockout HeLa cells. a,** Genotyping of *nup35* knockout clonal HeLa cell line. The protospacer adjacent motif (PAM) region is shown in orange, and the CRISPR-Cas9 nuclease target sequence in blue. Gene editing events are indicated in red at the bottom of the wild-type sequence. **b,** Expression of nucleoporins in Nup35 knockout cells. Western blot analysis of Nup93 subcomplex proteins as well as TNPO3, Nup153, and Nup358 from Nup35 knockout HeLa cell lysates. **c,** Genotyping of *nup153* knockout clonal HeLa cell line. The PAM region is shown in orange, and the CRISPR-Cas9 nuclease target sequence in blue. Gene editing events are indicated in red at the bottom of the wild-type sequence.

**Extended Data Figure 5: Knockdowns of CypA-dependent nucleoporins have minor effects on MLV infection.** HeLa cells were either single-knocked down (Nup35, POM121, Nup153, or CypA) or double-knocked down (CypA with FG-Nups, Nup35, POM121 or Nup153) for 48 h. The cells were infected with VSV-G-pseudotyped MLV and infected RFP-positive cells were counted by FACS. This analysis was performed in parallel to the experiments shown in Fig. 2d. Error bars show standard deviations of duplicates, representative of three independent experiments. Western blotting confirmed knockdown efficiency.

**Extended Data Figure 6: Nup35 knockdown impairs HIV-1 nuclear entry.** HeLa cells were transfected with control siRNA or siRNA against Nup35 for 48 h, then infected with increasing amounts of VSV-G-pseudotyped viruses in the presence or absence of 2.5 μM CsA. Firefly luciferase activity and percentage of RFP-positive cells (MLV infection) were measured at 48 h. Western blotting confirmed knockdown efficiency. This analysis was performed in parallel the experiments shown in Fig. 3b.

**Extended Data Figure 7: Subcellular distribution of soluble factors. a-c,** Distribution of CPSF6 and CypA in MT4 (**a**), Jurkat (**b**), and H9 (**c**) cells. Localization of CPSF6 (green), CypA (red), and lamin B1 (blue) in a Z-section of immunostained cells imaged by deconvolution microscopy. **d,** Nup35 depletion does not alter CypA subcellular localization. Localization of CypA (green) and lamin B1 (red) in a Z-section of immunostained HeLa cells imaged by deconvolution microscopy. **e,** Nup35 depletion does not alter CPSF6 subcellular localization. Localization of CPSF6 (green) and lamin B1 (red) in a Z-section of immunostained HeLa cells imaged by deconvolution microscopy.

**Extended Data Figure 8: C-terminal domain of POM121 is required for restriction of HIV-1 infection**. HeLa cells stably transduced with HA-tagged rhTrim5a/POM121 fusion constructs were infected with VSV-G-pseudotyped HIV-RFP or CA mutants (N74D or P90A) viruses, and percentage of RFP-positive cells was measured at 48 h.

**Extended Data Figure 9: Efficient HIV-1 infection in cells depleted of KPNB1 or Importin-7**. HeLa cells were transfected with control siRNA or siRNAs against TNPO3, POM121, KPNB1, or Importin 7 for 48 h, then infected with VSV-G-pseudotyped HIV-RFP or MX-RFP. After 48 h, RFP-positive cells were enumerated by FACS. Error bars show standard deviations of duplicates, representative of two independent experiments. Western blotting confirmed knockdown efficiency.

