## Supplemental files for "The HIV-1 capsid core is an opportunistic nuclear import receptor"

### Extended Data Figure 1

**a**

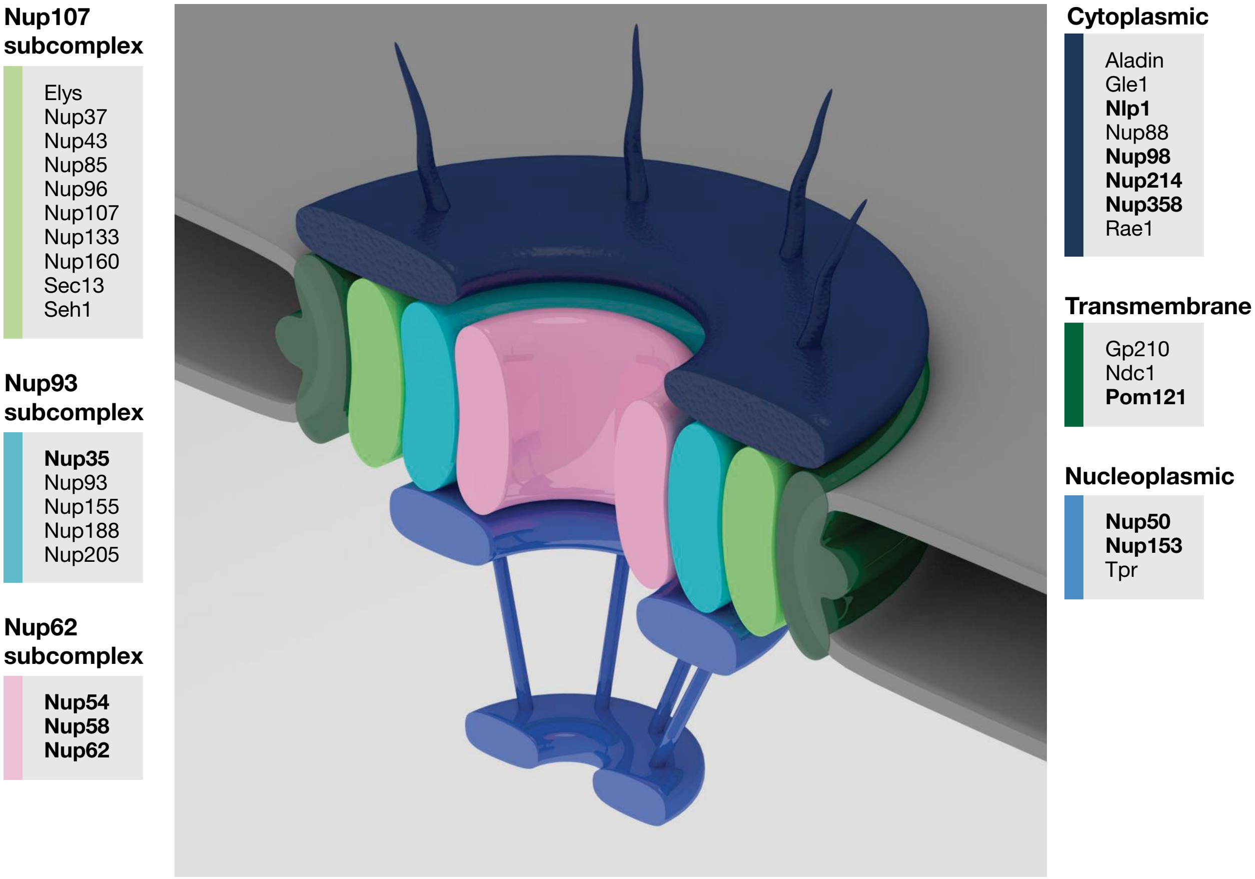


**b**

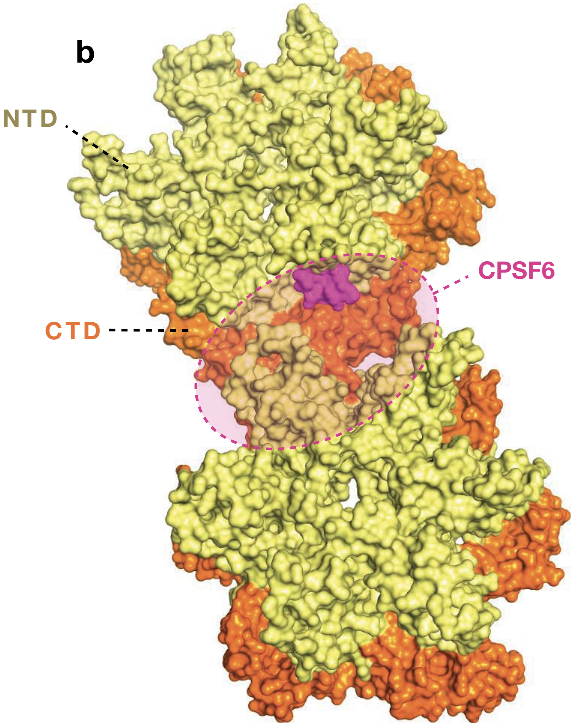


**c**

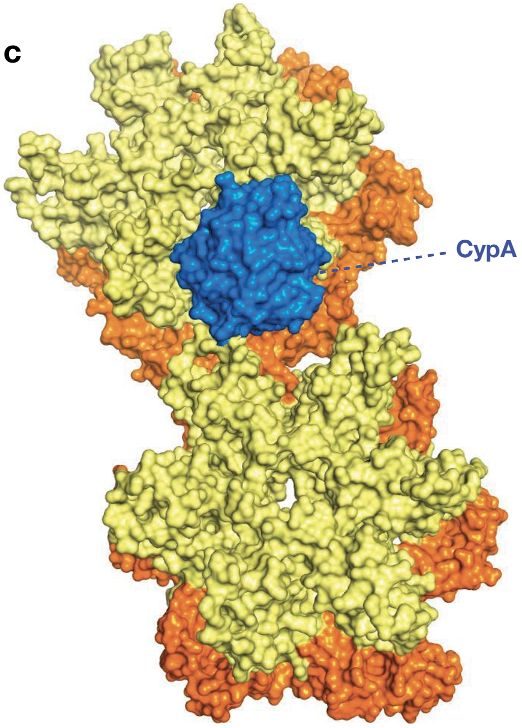

### Extended Data Figure 2

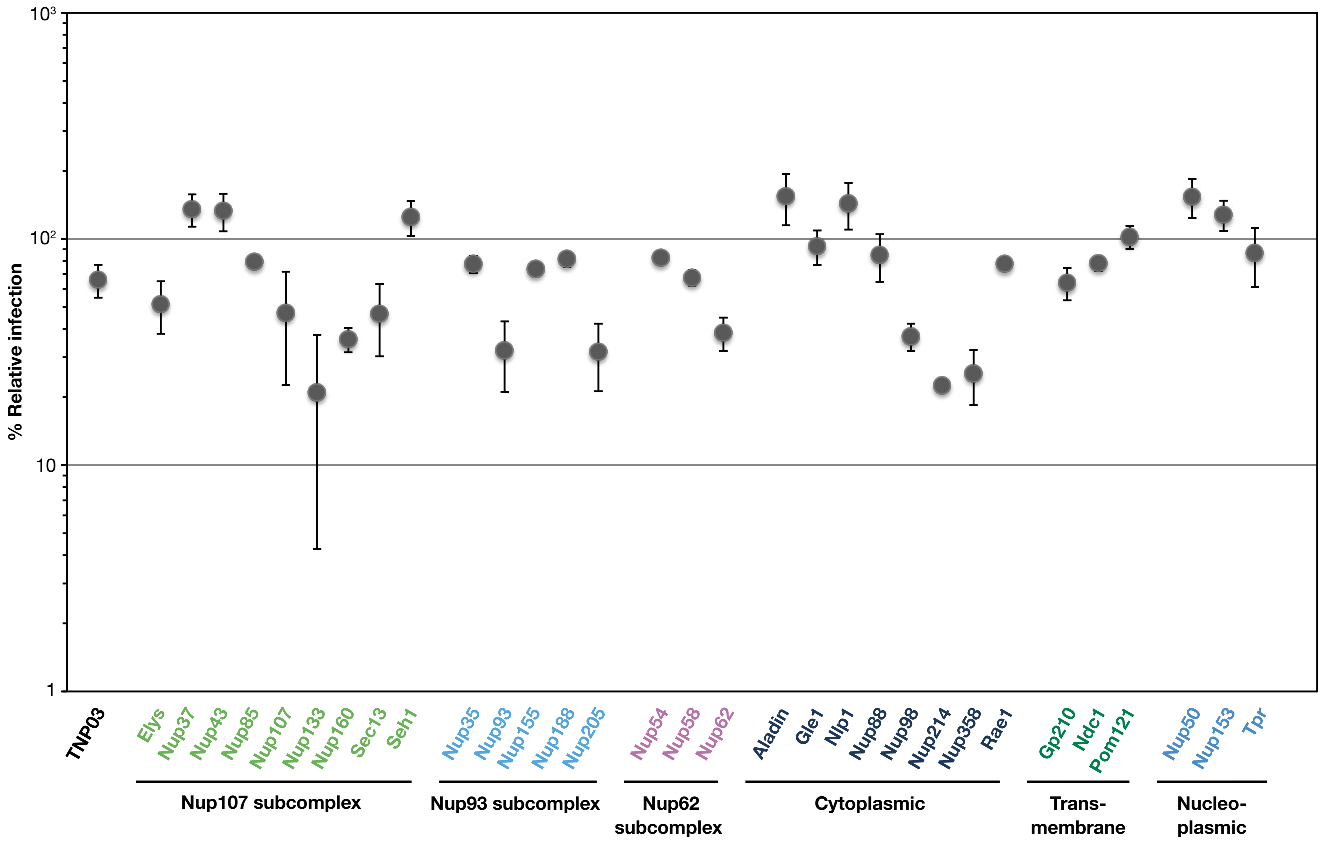

### Extended Data Figure 3

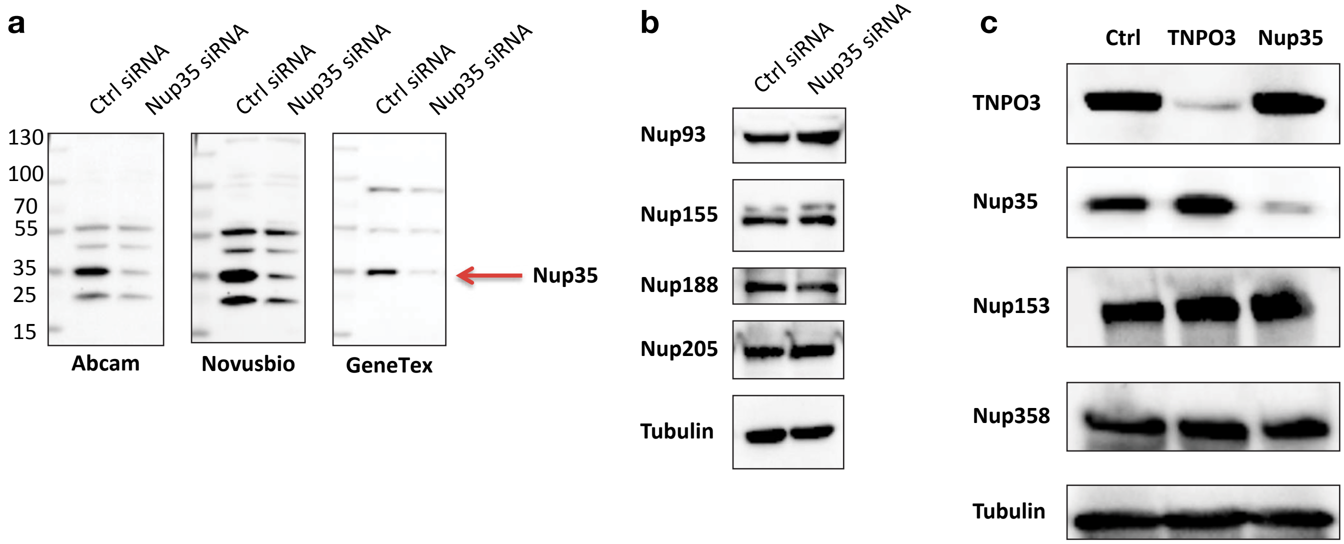

### Extended Data Figure 4

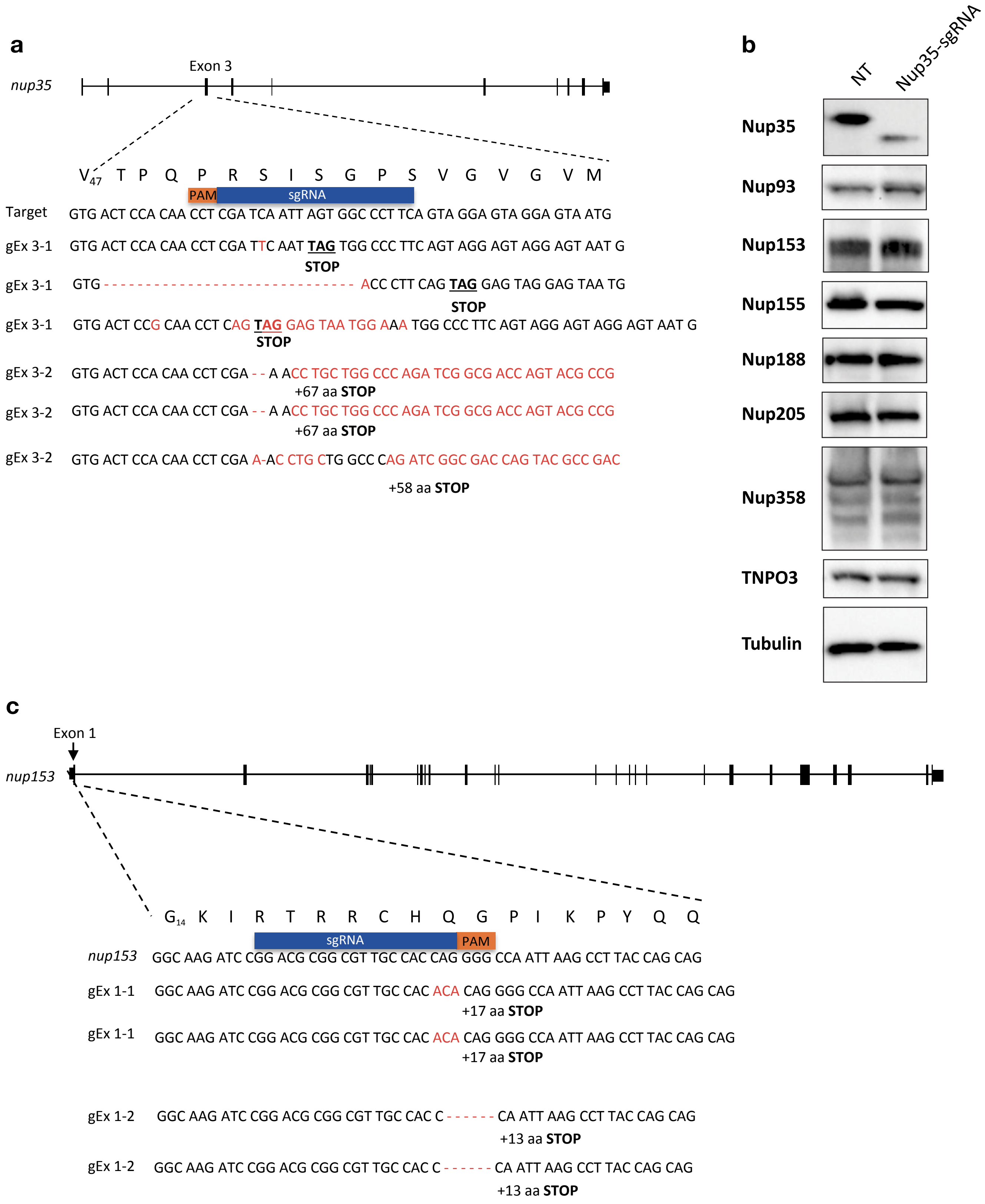

### Extended Data Figure 5

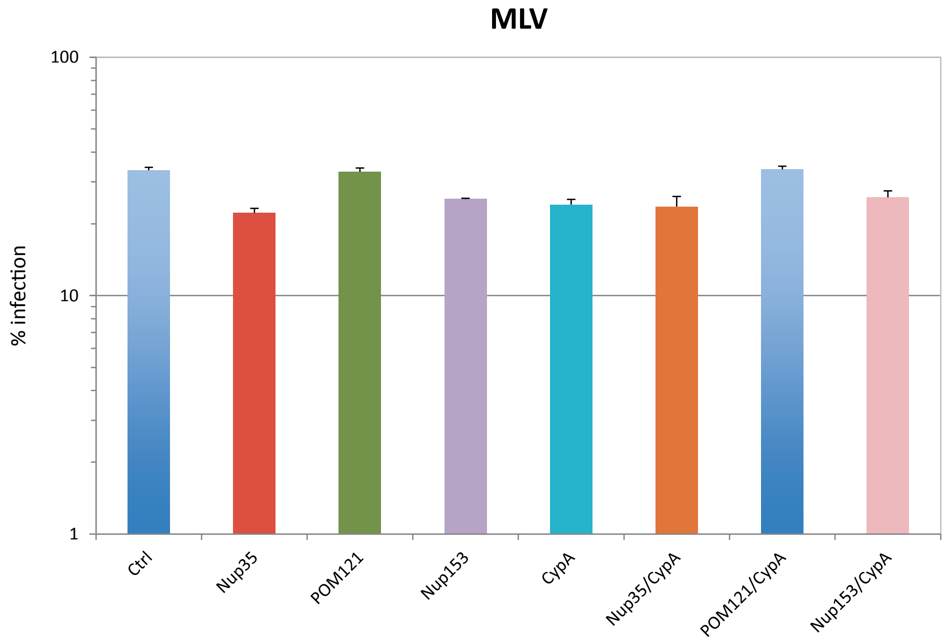


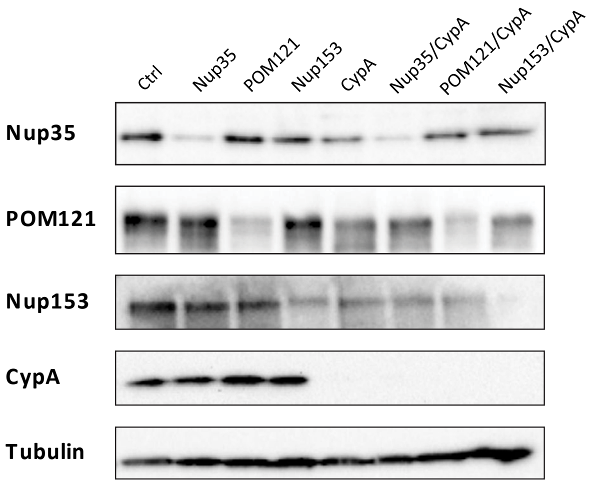

### Extended Data Figure 6

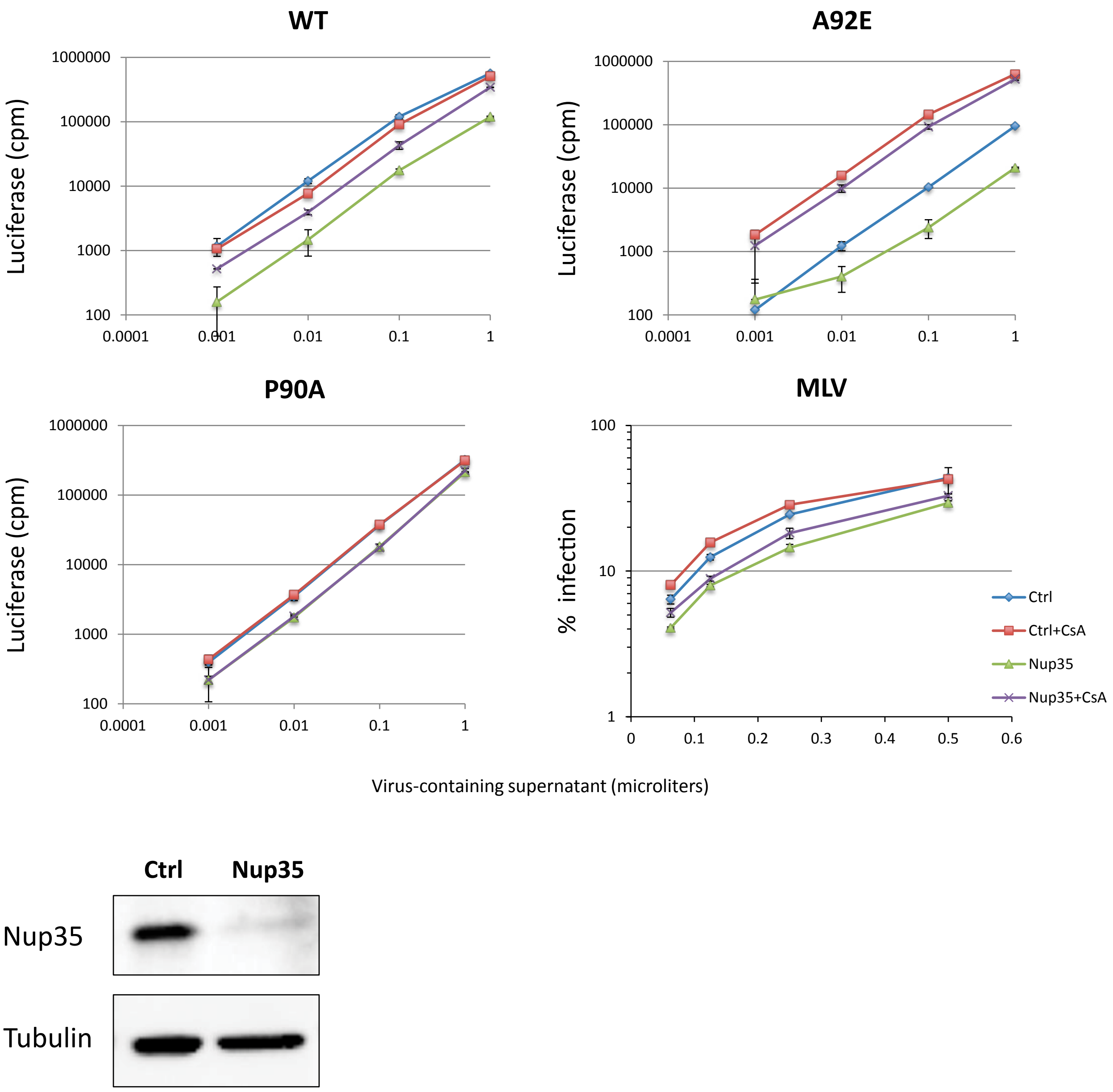

### Extended Data Figure 7

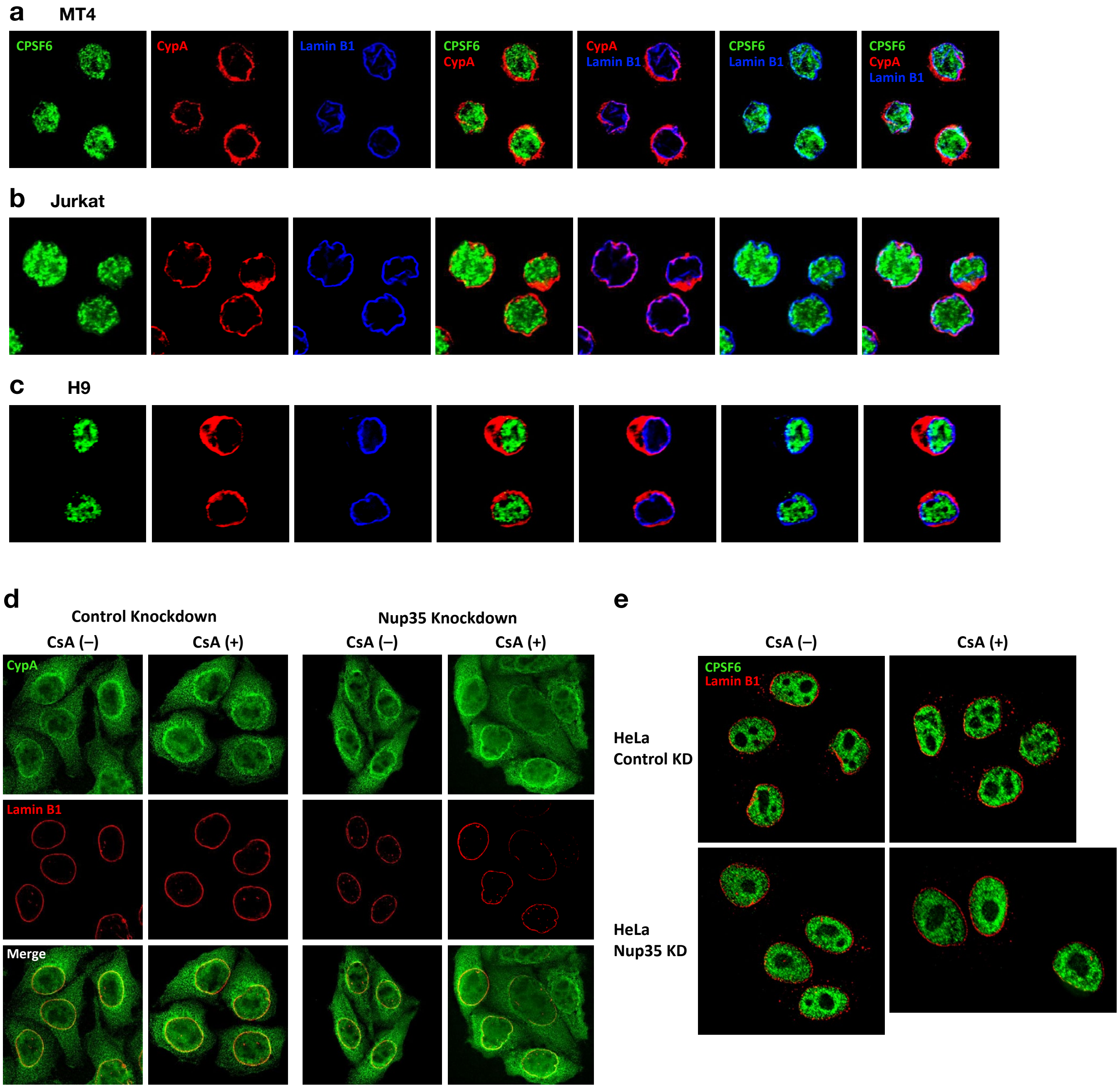

### Extended Data Figure 8

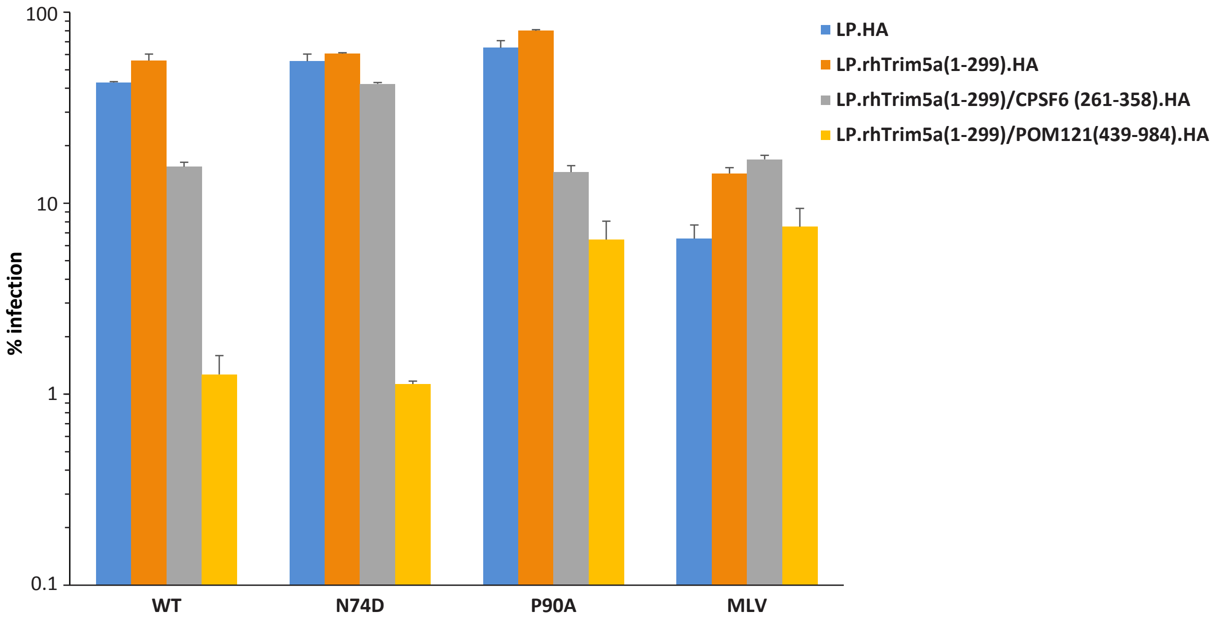

### Extended Data Figure 9

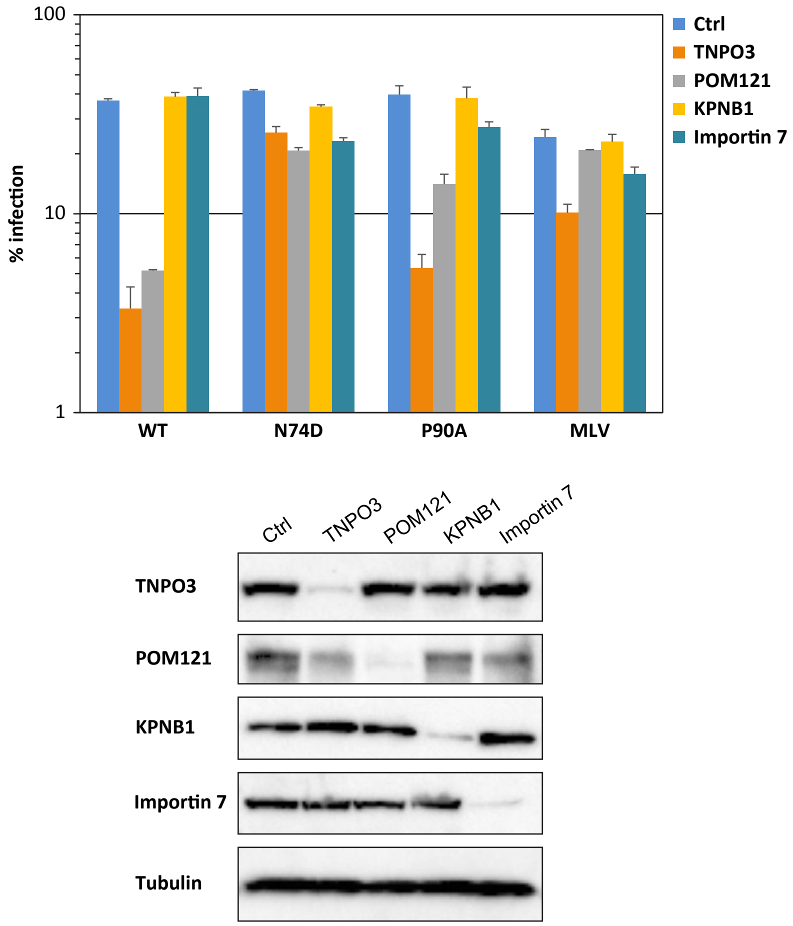
